# Dynamics of mRNA fate during light stress and recovery: from transcription to stability and translation

**DOI:** 10.1101/2022.06.10.495589

**Authors:** Aaron B. Smith, Diep R. Ganguly, Marten Moore, Andrew F. Bowerman, Yoshika Janapala, Nikolay Shirokikh, Barry J. Pogson, Peter A Crisp

**Affiliations:** Research School of Biology, The Australian National University, Canberra, ACT, 2601, Australia; CSIRO Synthetic Biology Future Science Platform, Canberra, ACT, 2601, Australia; Monash Biomedicine Discovery Institute, Department of Biochemistry and Molecular Biology, Monash University, Clayton, VIC, 3800, Australia; The John Curtin School of Medical Research, The Australian National University, Canberra, ACT, 2601, Australia; Australian Research Council Training Centre For Future Crops, Canberra, ACT, 2601, Australia; School of Agriculture and Food Sciences, The University of Queensland, Brisbane, QLD, 4072, Australia

**Keywords:** mRNA stability, translation, light stress, recovery, *Arabidopsis thaliana*

## Abstract

Transcript stability is an important determinant of its abundance and, consequently, translation. However, it is unclear the extent to which it is modulated between environmental conditions. We previously hypothesised that recovery-induced transcript destabilisation facilitated a phenomenon of rapid recovery gene down-regulation (RRGD) in *Arabidopsis thaliana* following stress, based on mathematical calculations to account for ongoing transcription. Here, we test this hypothesis, and investigate processes regulating transcript abundance and fate, by quantifying changes in transcription, stability, and translation before, during, and after light stress. We adapt syringe infiltration to apply a transcriptional inhibitor to soil-grown plants in combination with stress. Compared to measurements in juvenile plants and cell culture, we find reduced stability in a range of transcripts. We also observe transcript destabilisation during light stress, followed by stabilisation upon recovery. Alongside fast transcriptional shut-off in recovery, this destabilisation appears to facilitate RRGD. Translation was dynamic over the course of light stress and recovery, with substantial transcript-specific increases in ribosome-association, independent of changes in total transcript abundance, observed after 30 minutes of light stress. Taken together, we provide evidence for the combinatorial regulation of transcription, stability, and translation that occurs to facilitate responses to light stress and recovery.

## Introduction

Responses to stress have been the subject of much research; however, the post-stress recovery phase has been understudied and is likely to be equally complex and dynamic. Often, the molecular changes that occur at the onset of stress are protective in nature and divert resources away from growth and reproduction. Recovery, therefore, serves the purpose of resetting these processes in order to maximise growth potential. A balance is required between such resetting and maintaining the expression of acclimatory proteins, as a form of ‘memory’ (1, 2). Nonetheless, it may be counter-productive to maintain protective mechanisms indefinitely, as demonstrated by the growth penalty incurred by some stress tolerant plant lines (3). For instance, activity of the K^+^ efflux antiporter 3 (KEA3) was found to be important for relaxation of the energy-dependent portion of non-photochemical quenching (qE) following light stress (4). Increasing *KEA3* expression resulted in faster qE relaxation resulting in enhanced photosystem II efficiency under fluctuating light (5), which can influence CO_2_ fixation and biomass accumulation (6). This example highlights the potential in optimising stress recovery in food crops.

Transcriptome resetting has been characterised during recovery from multiple stressors. For example, sulphur starvation in Arabidopsis led to the up-regulation of many sulphur metabolism-associated genes, which returned to pre-stress levels within three hours of resupply (7). Recovery of rice to phosphate starvation occurred within one day of resupply, although full recovery took 31 days in line with re-establishing original phosphate content (8). On the other hand, rice recovered from submergence within hours (9). We previously highlighted that the Arabidopsis transcriptome is rapidly reset following light stress, whereby stress-induced mRNAs are down-regulated to pre-stress levels within 30 minutes; a phenomenon termed RRGD (10). Of note was the observation that RRGD loci exhibited far shorter half-lives during recovery compared to steady state measurements (11), suggesting transcript-specific destabilisation. The ability to rapidly alter the composition of the transcriptome gains particular importance during stress, when factors such as dehydration, heat, and oxidative stress can cause extensive cellular damage. While transcription can be adjusted to produce stress-associated transcripts, post-transcriptional changes, including modulating mRNA stability, are likely to permit faster changes in abundance; a valuable feature when a rapid response is required (12). Subtle shifts in stability can also greatly affect the ultimate number of mRNA molecules per cell over time (13). It is worth noting that recovery is more than just resumption of pre-stress gene regulation, as a diversity of recovery-induced gene expression programs are evident (10, 14). For example, the down-regulation of stress-induced transcripts during recovery is accompanied by the up-regulation of distinct genes, the products of which may play roles in recovery. This includes genes encoding RNA decay factors, such as the deadenylase-encoding *CAF1* (10). Whether these transcripts are translated proportional to their up-regulation is unclear, but it may indicate a rapid means of re-establishing pre-stress cellular conditions.

There are increasing observations of changes in mRNA stability during plant stress responses. Under high salinity, N^6^-methyladenosine promotes stabilisation of transcripts encoding salinity-associated proteins (15). During cold stress, the interplay between changes in DNA transcription and mRNA stability and transcription culminate into faster changes in overall expression of cold-responsive genes (16). Transcript degradation during stress appears to utilise the 5′ → 3′ RNA decay pathway through the activity of decapping proteins (DCP1, DPC2, DCP5, and VARICOSE) and EXORIBONUCLEASE (XRN) 4. For example, 5′ → 3′ RNA decay has been implicated in thermal acclimation (17, 18), photomorphogenesis (19), pathogen resistance (20), ABA signalling (21), and dehydration or osmotic stress (22, 23). Inhibition of the nuclear XRNs, resulting in RNA polymerase II read-through into stress-associated genes, has also been associated with drought tolerance (24, 25). These studies highlight the biological importance and potential specificity of the RNA decay machinery.

Translation is intimately tied with RNA decay in both cooperative and antagonistic ways (26). Competition for the 5′ cap can be a key determinant of mRNA stability, and high ribosome density and translation speed can be an indicator of elevated stability (27–29). Conversely, co-translational decay mechanisms are utilised to degrade transcripts on which ribosomes are stalled or elongating at a slowed rate, thereby permitting ribosome recycling (30, 31). Previously, evidence of co-translational decay in Arabidopsis plants undergoing light stress and recovery suggested that light-induced transcripts could be degraded while still polysome-bound, bypassing the need for dissociation from ribosomes (10). During heat stress, for example, a dramatic increase in monosomal loading relative to polysome loading is observed, which dissipates during recovery (32).

Preferential translation may permit a cell to make rapid changes in protein levels independently of transcription (33). As prioritisation of energy and resources is often paramount during adverse conditions, translation is often repressed globally with the exception of a subset of transcripts required for stress responses (34, 35). The repression of translation can involve degradation or transcriptional repression of genes encoding ribosomal proteins or translational factors (36, 37); however, a more rapid response is facilitated through dissociation of transcripts from initiation factors and ribosomes (38, 39). In many cases, the disassembly of polysomes is associated with formation of cytoplasmic foci called stress granules, which store complexes of translationally inert mRNAs, translation initiation factors, and other RNA-binding proteins thereby preventing their shuttling to, and degradation at, processing bodies (40). Upon removal of the stress, transcriptional up-regulation of the translation machinery occurs presumably in order to resume pre-stress protein production (14). This may be aided by the release of transcripts encoding ribosomal proteins from stress granules. For example, Arabidopsis heat shock proteins regulate the disassembly of stress granules following heat stress, allowing stored mRNAs encoding initiation factor complexes to commence translation (32).

We hypothesise that mRNA stability and translation are combinatorially modulated during recovery from light stress, which contributes to the resetting of protective mechanisms by shaping the cellular RNA pool available for translation. Previously, we were not able to assess whether the stability of individual transcripts shifted between unstressed, stressed, and recovery using conventional methods. An ongoing challenge with examining mRNA stability during stress is delineating changes in stability from changes in transcription. Accurate measurements of RNA decay requires either cessation of transcription through the use of transcriptional inhibitors, or *in vivo* labelling of mRNAs. The treatment of plants undergoing stress is limited by the nature and duration of the stress being examined. For example, delivery of RNA analogs via root feeding requires at least one hour (41); unsuitable for short term, transient stresses such as high light in which the transcriptome changes within minutes (42). Previous attempts to model RNA half-life in the absence of these treatments has had some success, but was heavily limited to up-regulated transcripts that displayed stepwise reductions in abundance (10).

To address the aforementioned shortcomings in prior reports, we established a method to perform transcriptional inhibition in leaves of mature soil-grown plants using syringe infiltration of the transcriptional inhibitor, cordycepin. This enabled us to compare genome-wide changes in mRNA stability between stress and recovery conditions *in situ*. We paired these with measurements of precursor mRNA (pre-mRNA) levels, as a proxy for transcriptional changes; and with translation rates, by profiling ribosome-bound mRNA during light stress and recovery. This combinatorial strategy revealed that RRGD was facilitated by interplay between regulation of transcription and mRNA stability. Whilst total mRNA abundance tracked with polysome-bound RNA levels (i.e. translation), this relationship was uncoupled during late light stress and early recovery, suggesting the occurrence of translational re-organisation.

## Results

### Locus-specific changes in transcription are evident during light stress and recovery

To determine whether transcriptional changes contribute to RRGD, we quantified levels of pre-mRNA, relative to mRNA, during 60 minutes of high light (HL) followed by 30 minutes of recovery (REC) for the RRGD loci (10) *HSP101* (*AT1G74310*), *ROF1* (*AT3G25230*), and *GOLS1* (*AT2G47180*) (Figure S1). In general, changes in pre-mRNA mirrored those in mature mRNA during HL and REC. However, for both *HSP101* and *ROF1*, pre-mRNA levels during REC dropped below pre-stress levels, indicating a major decrease in mRNA production. During the same period, mRNA levels remained elevated over pre-stress levels by 4-fold (*ROF1*) and 8-fold (*HSP101*). In contrast, *GOLS1* pre-mRNA levels followed mRNA levels more closely during REC and never dropped below pre-stress levels, despite mRNA levels dropping even more rapidly than in *HSP101* and *ROF1*. For all three genes, pre-mRNA levels remained above pre-stress levels in samples that experienced HL beyond 60 minutes, indicating ongoing transcription. These results suggest that locus-specific transcriptional changes contribute towards RRGD.

### Syringe infiltration of cordycepin effectively inhibits transcription in mature leaves

We first sought to quantify *in planta* transcript half-lives in mature soil-grown plants during stress and recovery. Owing to the limitations of existing methods, this is typically performed on juvenile seedlings, grown on nutrient-rich media, or with cell culture (11, 41, 43, 44). While these systems allow for effective administration of transcriptional inhibitors (e.g. cordycepin or actinomycin D) or uridine analogs for pulse labelling, they do not allow for studies of mature soil-grown plants alongside applications of external stimuli (e.g. abiotic stress). Therefore, it is currently unclear whether existing measurements of RNA half-lives are reflective of mature soil-grown plants. To address these limitations, we developed a method of administering cordycepin that allowed the simultaneous application of stress treatments. Syringe infiltration was considered a viable procedure for administering cordycepin into the cells of mature Arabidopsis leaves, which presented two advantages. First, it allowed for *in situ* treatment of soil-grown plants during light stress. Second, since the treatment was applied to individual leaves, a mock treatment could be applied to a separate leaf on the same plant as a control. We could also test for systemic effects by comparing these responses to a third untreated leaf.

We first evaluated the ability to inhibit transcription using syringe infiltration into individual leaves before exposure to light stress and measurement of light-responsive gene induction by qRT-PCR (Figure 1). While strong induction of *HSP101* and *HSP17.4B* was observed when infiltrated with a mock buffer, this was dramatically attenuated when infiltrated with cordycepin for 10 minutes (Figure 1 A-B). For example, *HSP101*, which was up-regulated 375-fold after 30 minutes of high light in mock treated leaves, was only elevated 4.5-fold in the presence of cordycepin. This equated to a 98.8% attenuation of the response. A similar block of induction was observed for *HSP17.4B*. We next examined how long cordycepin remained effective after infiltration into mature plant leaves; an important consideration for RNA half-life measurements. Leaves were infiltrated with cordycepin for a pre-incubation of 10, 30, or 60 minutes under unstressed (US) conditions, before exposure to 15 minutes of stress, with efficacy measured via induction of *HSP101* (Figure 1 C). Relative to the 32-fold induction in the absence of cordycepin, the highest induction (or lowest attenuation) achieved was approximately 8-fold when light stress was applied after 60 minutes of cordycepin incubation, equating to 75% inhibition. However, substantial transcriptional inhibition was observed after 10 (3.8-fold induction, 88% inhibition) and 30 minutes (1.5-fold induction, 95% inhibition) of incubation. It was uncertain whether the application of cordycepin to one leaf could cause systemic effects, which would affect the ability to mock infiltrate a separate leaf on the same plant as an internal control. We tested this by comparing gene induction in response to 15 minutes light stress in non-infiltrated and mock-treated leaves of separate plants subject to different treatments (Figure 1 D). These included untreated (U), mock infiltration (M), or mock infiltration on one leaf and cordycepin infiltration on a second leaf (M+C) on the same plant. Additionally, a non-infiltrated leaf was also taken from each plant for comparison. For non-infiltrated leaves, no significant difference in *HSP101* induction was observed between treatment regimes (ANOVA, P = 0.216), nor for mock treated leaves (unpaired Student’s t-test, P = 0.633). But in both cases, incremental increases in mean expression were observed as the number of infiltrated leaves per plant increased. While not statistically significant, this suggested that the infiltration itself could stimulate stress-responsive genes. Critically, it was clear that transcriptional inhibition following application of cordycepin did not impair the induction of genes profiled in distal untreated or mock-infiltrated leaves (compare to Figure 1C - 10 min). Taken together, we concluded that syringe infiltration presented an effective means of cordycepin administration to inhibit transcription in Arabidopsis leaves.

**Figure 1.**
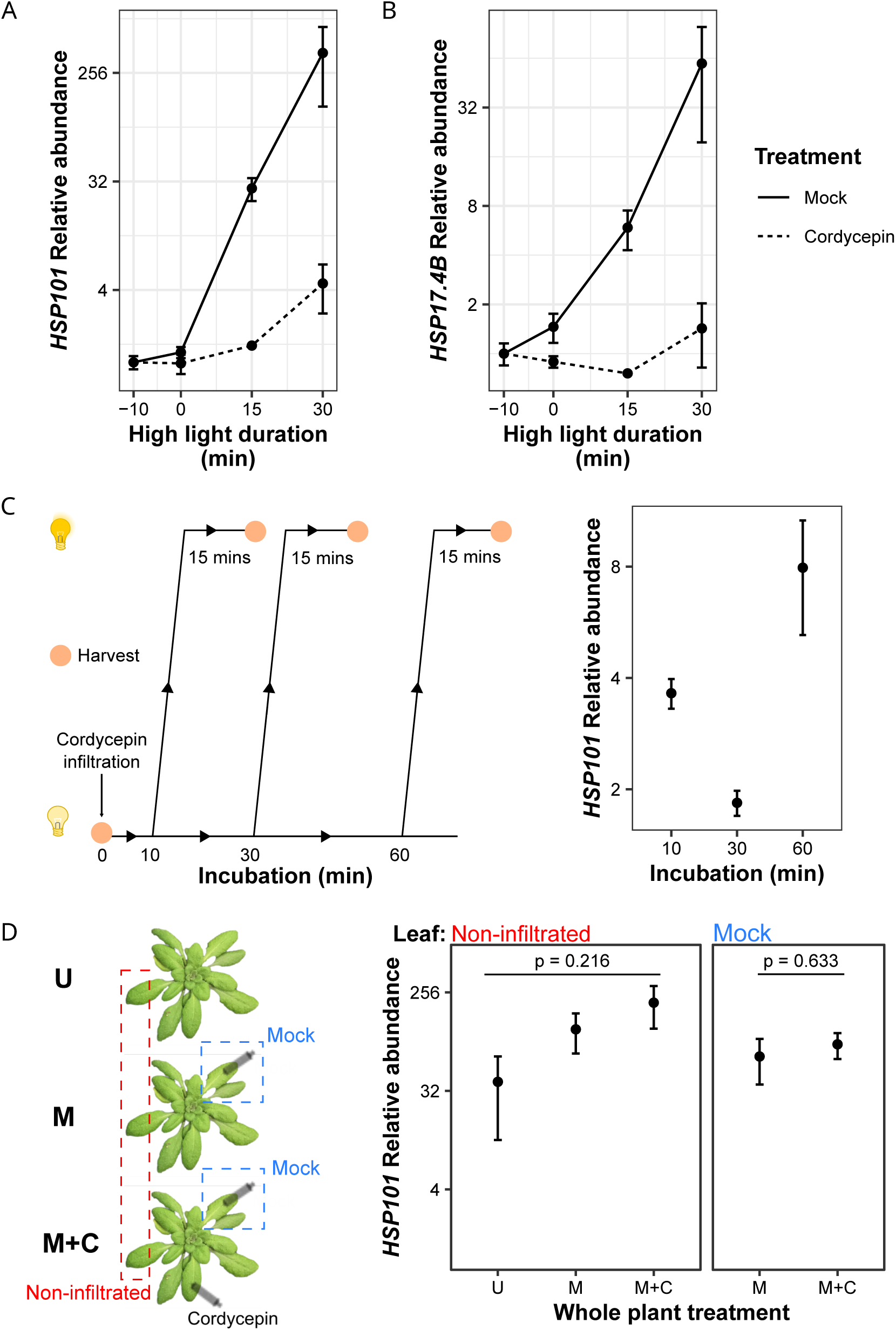
Syringe infiltration of cordycepin inhibits high-light induced transcription without systemic effects. (A-B) High light induction of *HSP101* and *HSP17.4B* was measured over 30 minutes folowing infiltration of individual leaves with mock or cordycepin solution. Points denote means, error bars denote standard error of the mean (n=2). (C) Arabidopsis plants were syringe-infiltrated with cordycepin and then incubated for 10, 30, and 60 minutes before exposure to 15 minutes of high light. (D) Arabidopsis plants were subject to three different infiltration regimes: untreated (U), mock infiltration (M), or mock and cordycepin infiltration (M+C), on independent leaves. Treatment was followed by 15 minutes of light stress prior to sampling untreated and mock infiltrated leaves. Points denote means, error bars denote standard error of the mean (n=3).

### Feedback regulation of mRNA stability in mature Arabidopsis leaves

With a functional cordycepin assay established for use under high light, a time course experiment was conducted to estimate RNA decay rates in US, HL, and REC conditions (Figure 2). In each instance, infiltration of mock or cordycepin solution was followed by at least three harvesting time points at 10 minute intervals (Figure 2 A). Based on the pre-incubation time course (Figure 1C) both 10 and 30 minutes were effective at inhibiting transcription. So, the minimal pre-incubation period of 10 minutes was used to allow for cordycepin to permeate cells to inhibit transcription effectively.

**Figure 2.**
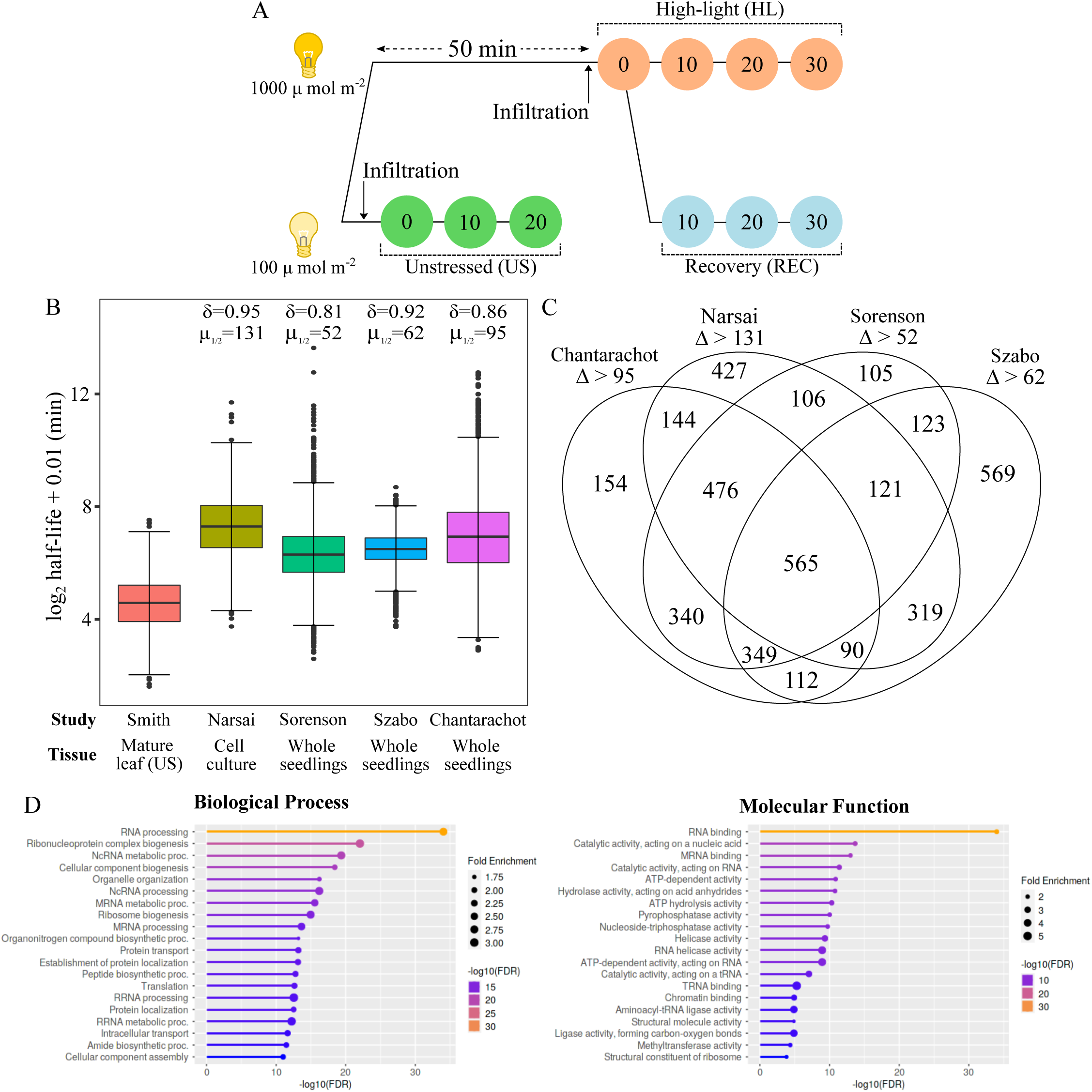
Sampling strategy to determine mRNA half-lives in Arabidopsis leaves. (A) Unstressed plants were harvested in standard growth conditions (US, 100 µmol m^-2^ s^-1^) at three time points in ten minutre intervals post-infiltration. High light and recovery plants were exposed to 60 minutes light stress (HL, 1000 µmol m^-2^ s^-1^), with infiltration occurring at 50 minutes. Following initial harvesting at 60 minutes HL, plants were either kept under HL or moved for recovery (REC, 100 µmol m^-2^ s^-1^) for three harvesting points at ten minute intervals. (B) Boxplots depicting log_2_ half-life determined for 4,497 genes under US conditions here, and their corresponding values from other studies. Box denotes median and interquartile range, bars denote 1.5xinterquartile range, and points denote outliers. δ denotes Cliff’s delta (the likelihood of observing a difference between groups) compared to this study. µ denotes the paired median difference compared to this study. (C) Venn diagrams representing the number of genes measured, in this study, with a reduced half-life greater than the median difference for each independent dataset assessed. (D) Top enriched GO terms for products of transcripts with reduced half-life in mature Arabidopsis leaves.

Infiltration of the two solutions was performed on independent leaves from the same plant to minimise biological variation between treatments, followed by mRNA sequencing. Multidimensional scaling plots highlighted similarity between replicates confirming the reproducibility of syringe infiltration and stress treatments (Figure S2). The primary source of sample differentiation was infiltration (Dim1), which was most pronounced after 20 or 30 minutes. The second source of differentiation was exposure to stress (Dim2), with HL and REC co-clustering separately from US samples. To examine whether syringe infiltration of cordycepin led to widespread transcriptional inhibition, differential gene expression was assessed between the first and last time points following cordycepin and mock infiltration (Figure S3, supplementary table 4). For mock treated samples, no particular bias toward up-or down-regulation was observed; by contrast, cordycepin treatment caused strong down-regulation in each condition (Figure S3 A). For example, during recovery 5,178 genes were down-regulated following cordycepin infiltration, while just 584 genes were upregulated. The up-regulation likely represents relatively stable transcripts that become comparatively more abundant as the RNA pool shrinks, leading to the appearance of increased expression. We utilised decay factor normalisation to reduce this artefact (44), however, some transcripts still showed a slight level of induction. Nonetheless, the induction of genes by the mock treatment was largely abolished in each condition (US: 95%, HL: 75%, REC: 74%) and the magnitude of down-regulation was greater following cordycepin treatment compared to mock. Taken together, this data confirms that widespread transcriptional inhibition was achieved in mature Arabidopsis leaves using syringe infiltration of cordycepin.

We next sought to calculate transcript half-lives across the transcriptome in unstressed Arabidopsis. Current measurements in Arabidopsis are determined using tissue culture with actinomycin D treatment (11), or nutrient-rich media-grown juvenile seedlings using uridine analogs (41) or cordycepin treatment (43, 44). We compared measurements with these systems to observations in mature soil-grown Arabidopsis leaves. Log-linear regression was performed using normalised fractional decreases of mRNA abundance in cordycepin infiltrated samples to determine *k_d_*, which allows for calculating a half-life that was used for comparisons. From the unstressed samples, half-lives could be computed for 6,711 genes based on statistical appropriateness of the fitted regression (supplementary table 5, p < 0.05). Half-lives were extracted from the aforementioned studies, for which 4,497 genes could be directly compared with our observations (Figure S3 B). This direct comparison hinted at shorter half-lives in mature leaf tissue compared to prior studies in seedlings and cell culture. Indeed, the transcript half-lives observed in this study had a median of 24.1 mins, which is substantially lower than the prior studies with median 79.2 - 157.6 mins (Figure 2 B). This was further supported by computing bootstrapped effect size parameters (mean difference, θ; median difference, µ_1/2_; Cliff’s delta, □) (Table 1). In particular, the bootstrapped median difference in transcript half-lives between datasets varied between 52 - 131 mins, with corresponding Cliff’s delta in the range of 0.81 - 0.95 (where a positive value represents the likelihood of observing a larger value). Larger values were observed when computing mean differences between datasets, likely caused by a small number of transcripts that displayed very large half-lives in some datasets.

**Table 1.** Effect size parameters for differences in transcript half-life between this study and other datasets

| Comparison study | Mean difference ($\theta$) | Median difference ($\mu_{1/2}$) | Cliff's delta (□) |
| --- | --- | --- | --- |
| Narsai et al. 2007 | 180 | 131 | 0.946 |
| Sorenson et al. 2018 | 94.8 | 52.4 | 0.808 |
| Szabo et al. 2020 | 70.1 | 62.1 | 0.922 |
| Chantarachot et al. 2020 | 352 | 94.8 | 0.858 |

Next, we explored which transcripts exhibited reduced stability in mature Arabidopsis leaves and their biological roles. To define transcripts with lower stability, we compared half-lives obtained here with each published dataset on a gene-by-gene basis. Those with lower half-lives, by a difference greater than the median difference, compared to each dataset were collated (Figure 2 C). There were 1,601 transcripts with consistently lower stability compared to the existing studies (lower than 3 out of 4 prior measurements; supplementary table 6). These had enrichments of GO terms relating to post-transcriptional and translational processes (Figure 2 D), such as RNA processing, ribosome biogenesis, RNA binding, and RNA catalysis (including pyrophosphatase and nucleoside triphosphatase activity). This suggests feedback regulation of RNA processing, splicing, and translation through altered mRNA stability in mature Arabidopsis leaves.

### Dynamics of transcript stability during light stress and recovery

Profiling transcript abundance in the presence of transcriptional inhibitors, under US, HL, and REC conditions, allowed for the exploration of broad patterns of decay and condition specific changes in stability (Figure 3). Hierarchical clustering revealed four patterns of decay, which we define as stable, slow, fast, and rapid (Figure 3 A). We explored whether any functional terms were enriched across these categories. Within the stable category, there were enrichments for organonitrogen compound (FDR = 2.1 x 10^-36^), amide (FDR = 2.4 x 10^-24^), and small molecule biosynthesis (FDR = 1.5 x 10^-29^), energy generation (FDR = 1.5 x 10^-29^), and response to cadmium (FDR = 4.2 x 10^-23^). Slow decay genes exhibited enrichments for organonitrogen compound biosynthesis (FDR = 7.6 x 10^-57^), including peptides (FDR = 5.6 x 10^-44^) and amides (FDR = 9.1 x 10^-43^), translation (FDR = 2.2 x 10^-44^), and RNA processing (FDR = 2.7 x 10^-42^). Genes within the fast category were enriched for chromosome (FDR = 7.4 x 10^-23^) and chromatin organisation (FDR = 3.9 x 10^-16^), organelle organisation (FDR = 2.7 x 10^-17^), RNA processing (FDR = 3.1 x 10^-12^), and negative regulators of gene expression (FDR = 2.7 x 10^-11^). Lastly, rapid decay genes encoded proteins involved in protein modifications (FDR = 3.7 x 10^-11^), especially ubiquitination (FDR = 2 x 10^-11^), and hormone signalling (FDR = 6.5 x 10^-6^).

**Figure 3.**
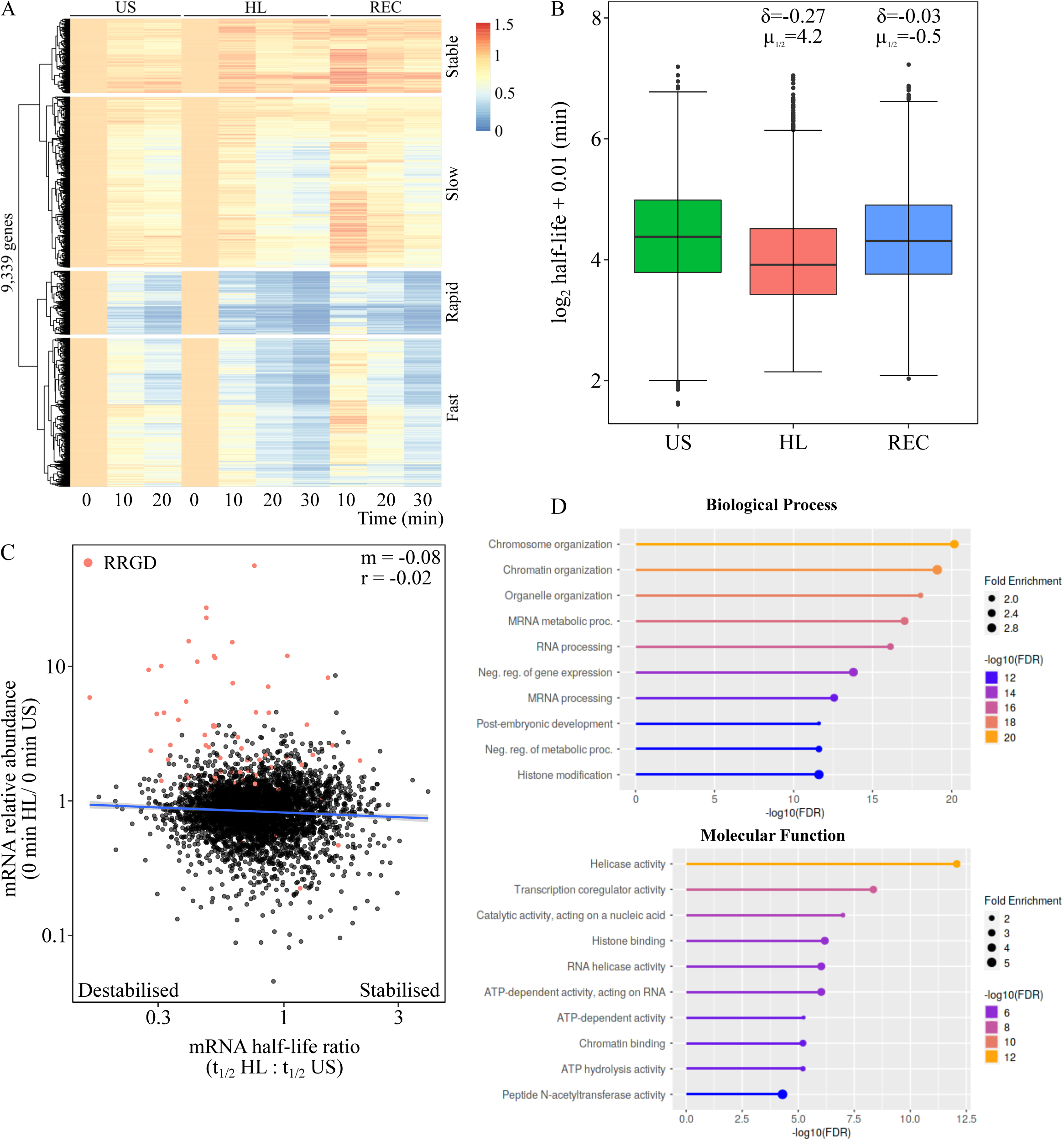
Transcript destablisation during light stress. (A) Heatmap with 1-dimensional hierarchical clustering depicting relative abundance of 9,339 genes in cordycepin-infiltrated samples under US, HL, and REC conditions. (B) Boxplots depicting log_2_ half-life determined for 3,960 genes under US, HL, and REC conditions. Box denotes median and interquartile range, bars denote 1.5 x interquartile range, and points denote outliers. δ denotes Cliff’s delta. (C) Scatter plot presenting the fold-change in mRNA abundance (0 min HL/0 min US), in mock treated-samples, versus the half-life ratio (HL/US) for 3,960 modelled transcripts. Points represent individual genes, line denotes fitted linear model, ‘m’ denotes regression coefficient, and ‘r’ denotes Pearson’s correlation coefficient. (D) Top enriched GO terms for products of transcripts destablised under HL.

Of particular interest was whether mRNA stability demonstrated conditionality. When comparing between conditions, transcript abundance appeared to decline at a greater magnitude under HL, suggesting faster degradation (or destabilisation) during stress. Curiously, there was a transient increase in abundance after 10 minutes REC, particularly in slow decaying transcripts. It is unclear whether this reflects residual transcription or is the result of slow decaying transcripts occupying a greater portion of the RNA pool as others are degraded. To quantify this destabilisation, we computed half-lives, for 3,960 genes that could be reliably modelled (p < 0.05), under US (Med t_1/2_ = 20.8 mins), HL (Med t_1/2_ = 15.1 mins), and REC (Med t_1/2_ = 19.8 mins, Figure 3B, supplementary table 7). While the bootstrapped median difference was only -1 min (□= -0.03) between US and REC, a difference of -4.21 min (□= -0.27) was observed under HL. This suggests that there is a bias towards destabilisation of transcripts under HL, which are re-stabilised during REC. We tested whether our RNA half-life analysis was biassed towards low abundance transcripts, which may be more prone to destabilisation. However, it was clear that the 3,960 transcripts captured in our analysis were of widely varying abundance (Figure S3 C). Comparing the half-life ratio under HL (t_1/2_ HL : t_1/2_ US) with the fold-change in mRNA abundance revealed a negligible relationship (Figure 3 C). However, there appeared to be a subset of highly induced transcripts that were destabilised and another portion of down-regulated transcripts that underwent stabilisation. Of the 388 RRGD loci defined previously (10), 70 were captured in this RNA half-life analysis. As expected, these were predominantly up-regulated (64/70 = 91.4%) when contrasted between HL and US in mock treated leaves. Indeed, these 64 genes comprised the majority of the most highly up-regulated genes by HL. Within this up-regulated subset, there was a clear bias towards destabilisation (55 / 64 = 85.9%), which suggests a coupling between induction (or increased abundance) and destabilisation. Conversely, these RRGD transcripts were largely down-regulated and stabilised in the transition from HL to REC (Figure S3 D).

We define 1,980 HL destabilised transcripts based on observing a reduction in half-life greater in magnitude than the median difference across transcripts between HL and US (i.e. 4.21 min, supplementary table 8). These genes were enriched for those encoding proteins involved in transcriptional regulation, such as chromosome and chromatin organisation, and transcription coregulator, histone binding, and helicase activities (Figure 3 D). Paired with this, were terms related to the (negative) regulation of metabolic processes and post-embryonic development. Once more, feedback regulation was observed, albeit to a lesser extent, with enrichments in mRNA metabolism and processing, and RNA helicase activity. Therefore, light stress appears to induce the destabilisation of transcripts involved in both transcriptional and post-transcriptional regulation, as well as the transition towards reproductive growth. Since it appeared that destabilisation of transcripts occurred under HL, we investigated the timing of this destabilisation (Figure S4). To test this, plants were exposed to high light for 60 minutes, with subsets of plants infiltrated with mock or cordycepin solution at 5, 20 and 50 minutes. Samples were then harvested at 20, 30 and 40 minutes to measure decay rates using qRT-PCR of *AT3G14200*, which exhibited HL-induced destabilisation (Figure S4 A). Plants infiltrated at 5 minutes HL exhibited an increase in *AT3G14200* in the 10 minutes following infiltration, which may be attributed to residual transcription and RNA processing while the cordycepin took effect on newly synthesising transcripts (45). However, transcript levels did decline from 20 to 40 minutes. In both the 20 minute and 50 minute treatment groups, steeper and more consistent declines were observed in the 30 minutes following infiltration. To quantify these differences, RNA half-life was modelled using the period 10 to 30 minutes post-infiltration. The half-life of *AT3G14200* after 5 minutes HL was estimated at 35.58 minutes, but this model was not significant (P_adj_ = 0.149). By contrast, the half-lives following infiltration at 20 and 50 minutes HL were estimated at 19.91 and 16.45 minutes, respectively (P_adj_ = 0.00027). This indicated that *AT3G14200* transcripts were more stable during the initial period of high light stress and were subsequently destabilised.

The finding that *AT3G14200* is destabilised based on the duration of stress suggests that RRGD, typically observed after 60 minutes, may not occur with shorter durations of HL. To test this, plants were exposed to 10, 30, 60 and 120 minutes HL before being moved to REC (Figure S4 B-D). Quantification of relative changes in canonical RRGD loci: *HSP101*, *ROF1*, and *GOLS1* using qRT-PCR revealed that 10 minutes HL did not cause recovery-induced down-regulation. By contrast, clear declines in transcript levels were observed when moved to REC after 30 and 60 minutes HL, while a small increase was observed followed by a decline after 120 minutes. Gene expression patterns seen in *GOLS1* were similar, though declines in transcript levels were observed in the last 10 minutes of REC after each HL length. Estimations of RNA half-lives were modelled for each REC period, however, following adjustments for multiple comparisons no model was significant (p < 0.05). Nonetheless, they did indicate the same general trend, with the most notable down-regulation being observed following 60 minutes HL. This data reinforced the cordycepin data obtained for *AT3G14200* and indicated that RRGD loci remain comparatively stable in the early stages of HL before being destabilised.

### Translational regulation occurs during late stress and early recovery

To investigate the interplay between transcript stability and translation during HL and REC, we sequenced polyribosome (polysome)-bound mRNAs, representing actively translated transcripts. This was done after 30 and 60 mins HL, and 7.5, 15, and 30 minutes REC (supplementary table 9-11). During polysome fractionation, we produced a spectral profile to identify specific fractions and test for shifts in the translational landscape (Figure S5). These profiles were consistent with recent work that optimised ribosome profiling from plant tissue using buffers of lower ionic strength (46). Indeed, we were able to distinguish monosome and polysome fractions, which separate based on the number of ribosomes bound to a transcript (Figure S5 A). The amount of RNA bound in each fraction was highly consistent between samples, indicating little change in transcript abundance loaded onto monosomes relative to polysomes (Figure S5 B). Although no major disruption was observed, individual transcripts may still undergo substantial translational regulation. To examine this, we sequenced the total and polysome-bound mRNA populations and calculated fold-changes between time points on a per-gene basis, which were then compared between the two populations (Figure 4). Multidimensional scaling was used to assess the similarity in polysome-bound and total mRNA quantification across samples, which confirmed reproducibility between biological replicates (Figure S5 C). Changes in total mRNA and polysome-bound mRNA were highly correlated (Pearson’s r = 0.81) in the early stages of stress (0-30 minutes) indicating loading of transcripts onto polysomes proportional to changes in transcript abundance (Figure 4 A). However, during the later period of stress (30-60 minutes) the two populations were substantially less correlated (Pearson’s r = 0.47). At early REC time points, the changes between the two populations were still weakly linked (Pearson’s r = 0.25, Figure S5 D). However, this correlation was gradually restored over 30 minutes REC (Figure 4 A, Pearson’s r = 0.77). This decoupling of total and polysome-associated mRNA abundance during late stress and early recovery is indicative of translational regulation on a subset of transcripts. Notably, this suggests that changes in translation are occurring independently of a concomitant change in total transcript abundance.

**Figure 4.**
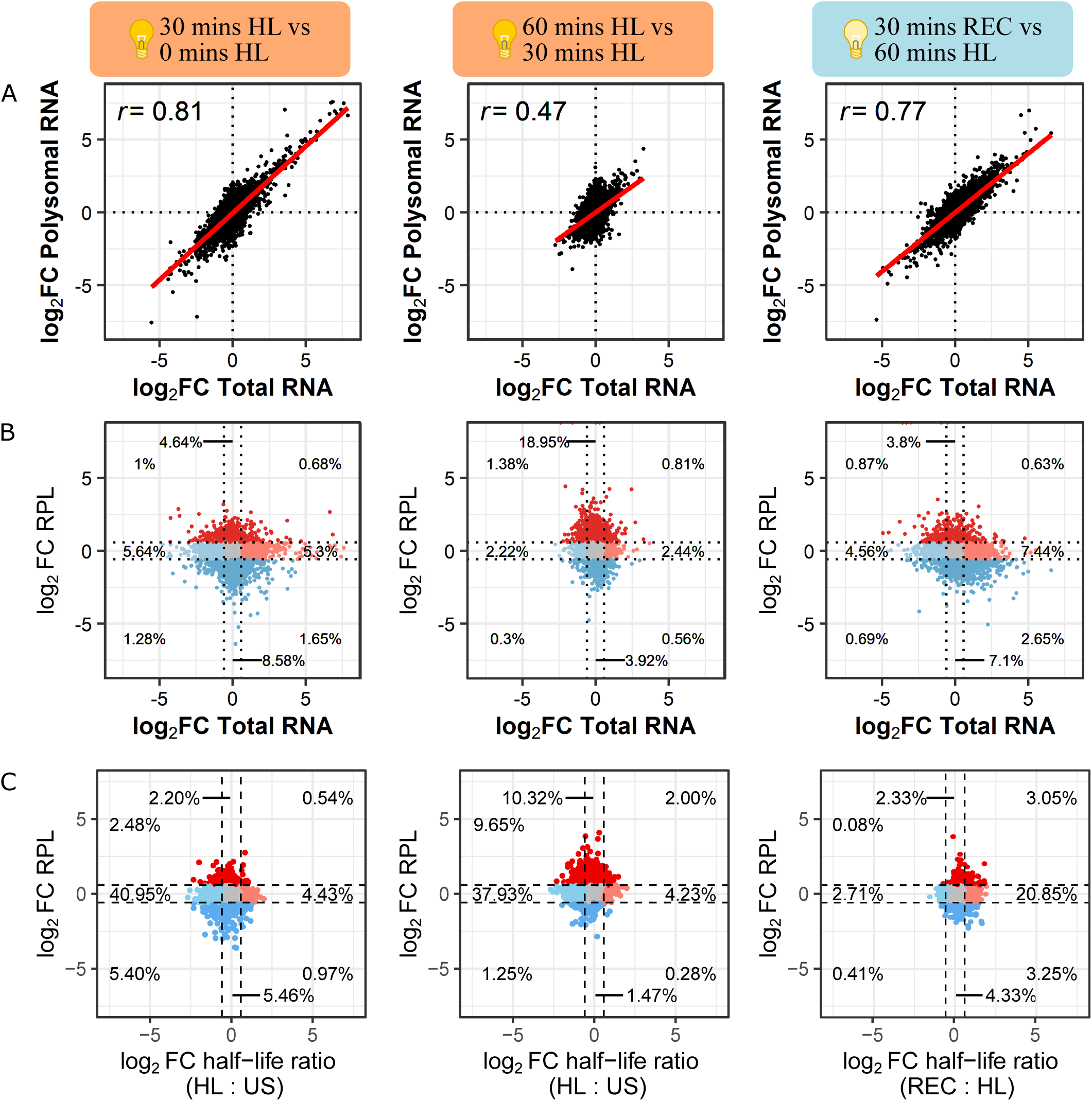
Translational regulation during light stress and recovery. (A) Scatter plots comparing fold-changes in total and polysome-bound mRNA populations between time points. (B) Scatter plots comparing fold-changes in total mRNA and relative polysome loading (RPL = polysomal mRNA / total mRNA) between time-points. (C) Scatter plots comparing fold-changes in RPL and half-life ratios between time-points. relating transcript half-life to their abundance in the polysome-bound mRNA fraction and RPL. Colouring indicates fold change greater (red) or less (blue) than 1.5 for either total mRNA or RPL. Numbers denote proportion of transcripts detected in each section. Points represent individual transcripts, line denotes fitted linear model, and ‘r’ denotes Pearson’s correlation coefficient.

As ribosomes are recruited onto transcripts across the total cellular pool, it can be informative to examine the ratio between polysome-bound and total mRNA. Since ribosome density was not measured, this is not a reflection of translation efficiency. Therefore, to avoid confusion, we refer to this as relative polysome loading (RPL), where a value of one indicates that polysome-associated and total mRNA occur at proportionally the same abundance. It is important to note that changes in RPL can occur via differential recruitment of ribosomes to transcripts (numerator) or changes in total transcript abundance (denominator). Notably, RPL can increase even if levels of polysome-associated mRNA decrease, provided this occurs at a slower rate than a reduction in total mRNA abundance. Therefore, comparing changes between RPL and total mRNA can be used to more easily identify shifts that occur in only one population. This comparison was performed for three periods: early HL (0-30 mins), late HL (30-60 mins), and REC (Figure 4 B). On a broad scale, clear differences could be observed between the three periods although a striking proportion of transcripts showed changes in RPL with no change in total mRNA abundance (early: 13.22%, late: 22.87%, REC:10.9%). In each period, these changes were enriched for regulation in opposing directions with early HL and REC being biassed towards reduced RPL whereas late HL showed a striking enrichment of genes with increased RPL. Such results indicate a bias toward translational repression at the onset of stress, followed by increased polysome loading. Furthermore, these 858 transcripts with increased RPL during late HL (supplementary table 12) were enriched for functions related to transcription (e.g. transcription regulator activity, FDR = 8.7 x 10^-5^), macromolecule methylation (e.g. RNA methylation, FDR = 6.3 x 10^-5^), RNA processing (FDR = 2.1 x 10^-8^), splicing (FDR = 1.2 x 10^-4^), and organelle organisation (FDR = 1.2 x 10^-4^). Notable too, were the changes in the proportion of transcripts showing differences in RPL relative to total mRNA, reflecting disproportionate changes in polysome-association compared to transcript abundance (Figure 4 B). During early stress, 20.8% and 19.2% of transcripts exhibited changes in RPL and total mRNA, respectively, a 1.1:1 ratio. During late stress, 25.9% and 7.7% of transcripts had changed RPL and total mRNA, respectively, a 3.4:1 ratio (3-fold increase). In recovery, this returned to 15.7% and 16.8% changes in RPL and total mRNA, respectively, reflecting a 0.9:1 ratio. Overall, these results highlight the extent of regulating translation during light stress and recovery, with the later stress period (30-60 minutes HL) being a dynamic period for translation showing increased ribosome-loading of many transcripts independently of changes in total mRNA.

Since transcript stability may influence translation, we examined the relationship between half-life and RPL during the same early HL, late HL, and REC periods. We hypothesised that destabilised transcripts would show reduced polysome-association. To do this, we related the fold-change in RPL, measured for each period, to the half-life ratio (i.e. the change in stability) observed under HL and REC for 3,905 genes detected in each dataset (Figure 4 C). There was no clear relationship between changes in RPL and mRNA stability. Under both periods of HL, there was a large proportion of transcripts that were destabilised yet had unchanged RPL indicating that the majority of destabilised transcripts retained similar levels of polysome recruitment. Conversely, during recovery there is a clear increase in the stability of many transcripts yet the majority of these also showed no difference in RPL. Taken together this suggests that changes in mRNA stability are not strongly linked with changes in polysome-association or dissociation.

### Transcript dynamics of RRGD genes likely impacts their translation

Previously, we defined RRGD genes as those up-regulated at least three-fold in mRNA abundance after 30 minutes HL. These were categorised based on their expression profiles: category 1 genes remained up-regulated through HL and REC, category 2 genes were down-regulated during REC, and category 3 genes were down-regulated after 30 minutes HL (10). However, it was unclear whether these dynamics had functional consequences. Therefore, we investigated whether the mRNA dynamics displayed by RRGD genes impacted translation. Examination of polysome-associated transcripts found that they exhibited an expression profile concordant with that observed in total mRNA (Figure S6). This suggests that, contrary to the decoupling observed on a global scale, the free and polysome-bound mRNA populations were tightly linked for RRGD genes. That is, changes in mRNA abundance of a RRGD gene generally led to a corresponding change in its translation. Furthermore, this indicates that the phenomenon of RRGD is likely to have tangible cellular consequences with respect to protein production.

Increased indices of co-translation decay were also observed in RRGD genes, suggesting its involvement in clearing these transcripts during recovery (10). Therefore, it was hypothesised that RRGD genes would exhibit reduced RPL as their polysome-associated transcripts are degraded at a higher rate than free unbound transcripts. To test this, changes in RPL were compared between each category of RRGD gene across our time course (Figure 5). Genes in categories 1 and 3 tended to have an increase in RPL during 30-60 minutes HL (Figure 5 A-C). This was particularly notable for category 3 genes that exhibited reduced total mRNA abundance (Figure 5 D-F). However, all categories of RRGD genes exhibited declines in RPL during REC, indicating either unloading (e.g. category 1) or degradation (e.g. categories 2 and 3) of polysome-associated transcripts.

**Figure 5.**
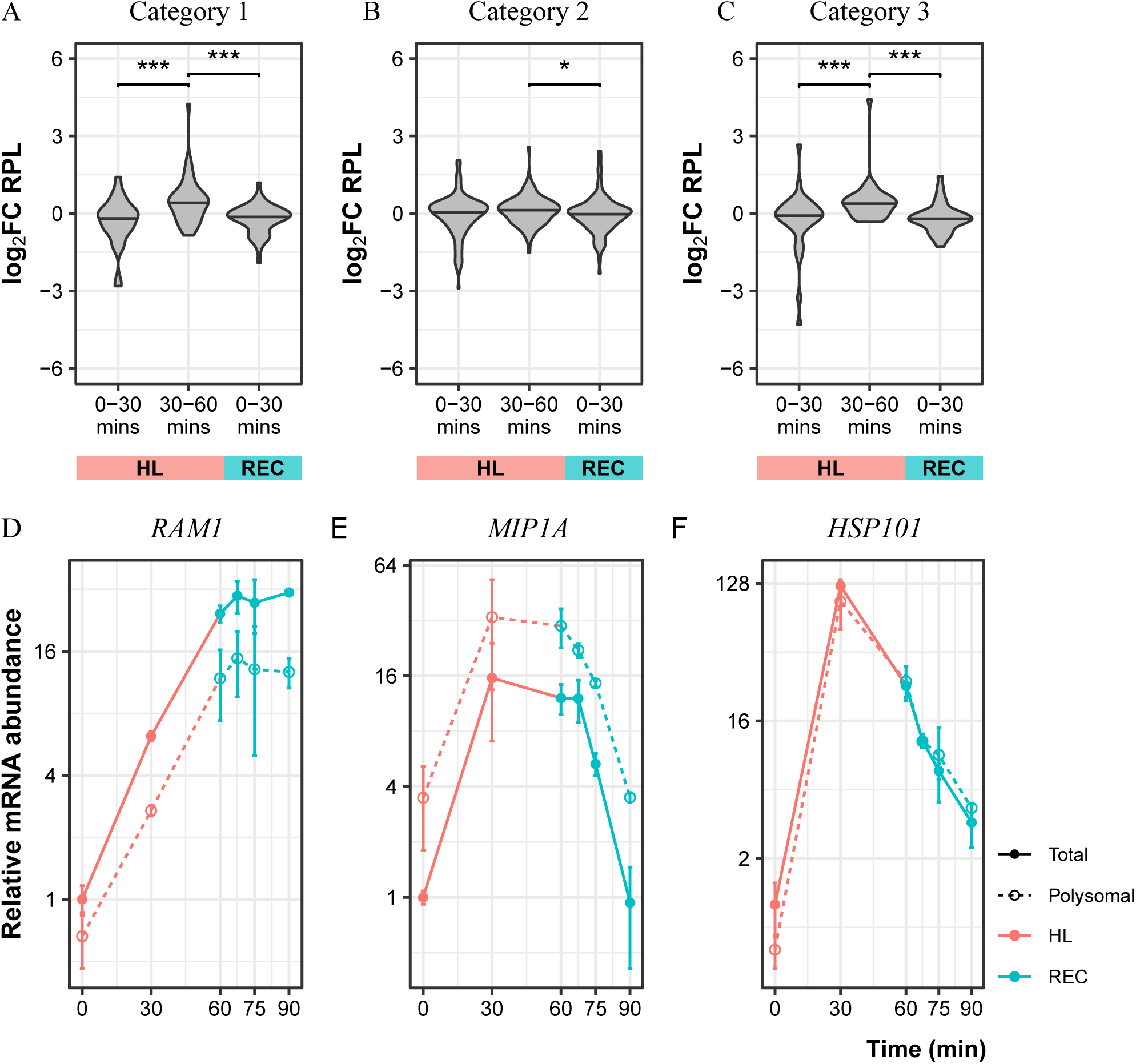
RRGD transcript dynamics impacts translation. (A-C) Light-induced genes were divided into categories based on expression profiles during late stress and recovery. Changes in RPL were compared across the three 30 minute time periods and are represented as violin plots. Median fold-change was compared using Wilcoxon Rank-sum test, * denotes P < 0.05, *** denotes P< 0.001. (D-F) Examples of gene expression profiles in each category. Points denote mean, error bars denote SE.

## Discussion

To regulate gene expression during stress and recovery, plants can alter transcription, mRNA stability, and/or translation. Here, we characterise changes in each during light stress and recovery, a highly dynamic period for mRNA abundance (10). Our focus was the extent to which mRNA stability was modulated to control RNA abundance, and the subsequent impacts on translation. Light-induced changes in transcription were based on quantifying changes in pre-mRNA levels for *HSP101*, *ROF1*, and *GOLS1*, all of which showed decreases upon perception of recovery. Meanwhile, mRNA destabilisation occurred during light stress followed by re-stabilisation during recovery. Based on detailed expression profiling of *AT3G14200*, this destabilisation appears to occur after 30 minutes of high light, at which point peak expression is reached. Similarly, recovery-induced down-regulation was not observed for *HSP101*, *ROF1*, and *GOLS1* in plants exposed to 5 minutes of stress, suggesting that destabilisation is dependent on the duration of exposure. Further, we found that light induced transcripts showed concordant changes in polysome loading and total mRNA levels. Based on our findings, we propose the following model of mRNA dynamics during light stress and recovery in Arabidopsis (Figure 6). Onset of stress is associated with a burst of light-induced transcription, which occurs for 30 minutes. Following this, mRNA stability decreases and transcription slows, leading to slower accumulation in overall abundance. Yet, preferential ribosome-association of specific transcripts, encoding light-induced and regulatory proteins (e.g. transcription and RNA processing), occurs during late stress. Upon entering recovery, transcription of stress induced mRNAs largely ceases, virtually shutting off for some genes. Due to the prior transcript destabilisation, a rapid down-regulation in total and polysome-associated mRNAs occur causing a resetting of the transcriptome to near pre-stress levels. In this way, changes in mRNA stability act as a precursor to RRGD, maximising the impact of the transcriptional shut-off by accelerating transcript turnover during this period.

**Figure 6.**
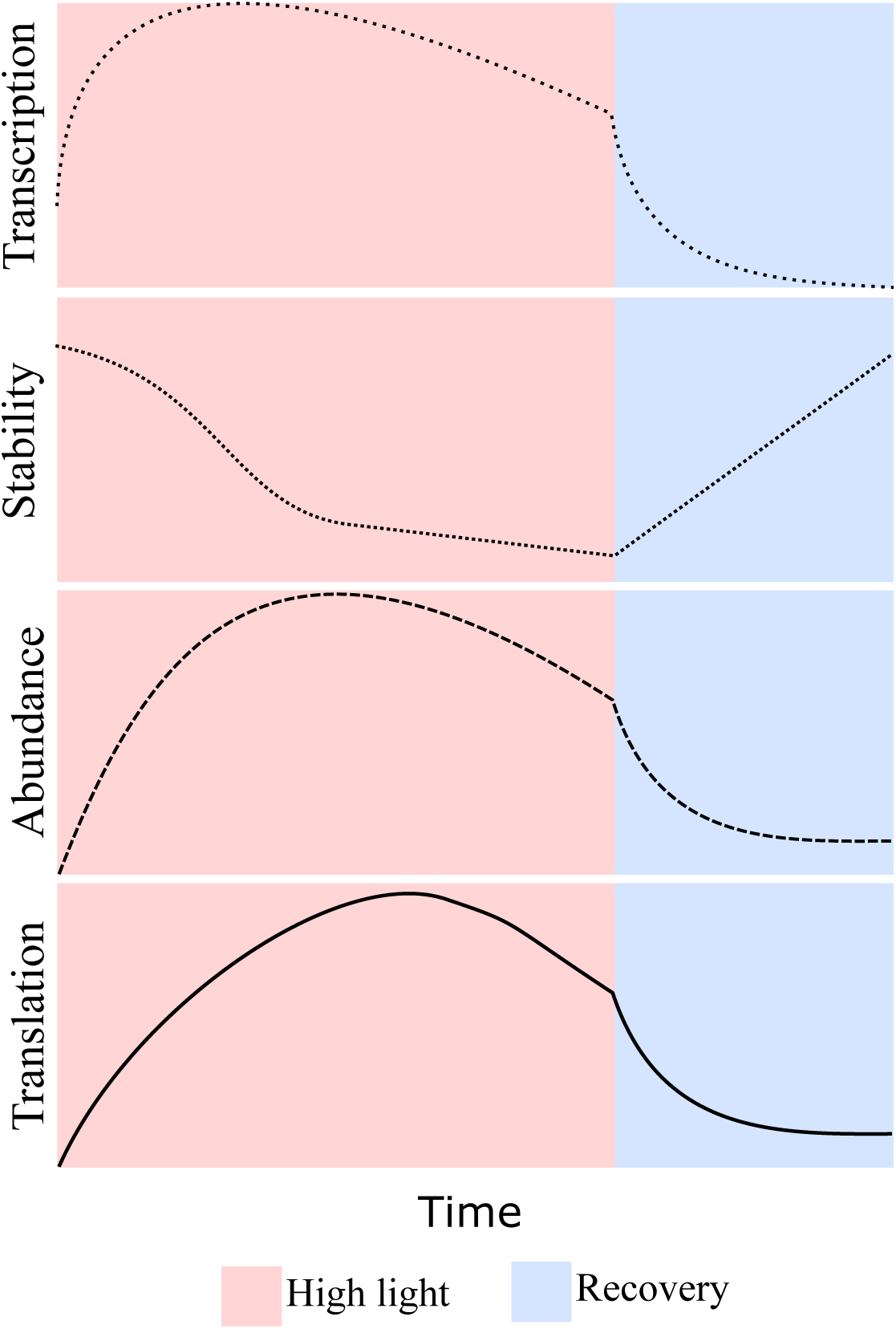
Stress-induced transcript destabilisation facilitates rapid recovery gene down-regulation. A proposed model of RRGD transcript dynamics during high-light and recovery. While transcription appears to rapidly decrease during recovery, the corresponding down-regulation in transcript abundance and resetting of the transcriptome is likely dependent on prior destabilisation.

### Dynamic mRNA stability during plant development and light stress

Half-lives of mRNA were measured in mature Arabidopsis leaves using a newly developed method, based on syringe infiltration of cordycepin. This allowed us to test for differences in mRNA stability in older plants, since existing measures have been performed in juvenile seedlings. Indeed, we observed a substantial decrease in half-life for the majority of genes that could be contrasted with existing datasets (11, 41, 43, 44). Those transcripts exhibiting reduced stability in mature leaves were strongly enriched to encode proteins involved in post-transcriptional processes, including RNA processing and catalysis, RNA binding, ribosome biogenesis, and translation (Figure 2 D). Contrary to a mature leaf, juvenile seedlings growing in nutrient-rich media will likely be undergoing higher rates of cell division (47–49) and chloroplast development (48, 50). When paired with a ready supply of the requisite macromolecules, this may be driving higher rates of protein synthesis and turnover (51, 52). This observation may, therefore, reflect autoregulation of post-transcriptional processes allowing greater responsivity (i.e. reduced mRNA stability allows for shorter transition times) to tailor protein synthesis.

We previously hypothesised that mRNA stability was modulated during recovery based on observing a rapid decline in the abundance of light-induced transcripts (10). The use of syringe infiltration allowed *in situ* application of cordycepin to plants undergoing stress and recovery. Contrary to expectations, we observed transcript destabilisation under high light followed by re-stabilisation upon recovery (Figure 3 B). Typically, half-life is measured over a longer period over hours rather than 30 minutes as performed here. Therefore, our analysis is likely biased for relatively unstable transcripts since longer time courses are needed to model statistically significant changes of more stable transcripts. A pronounced spike in relative abundance was observed for a subset of transcripts after 10 minutes of recovery in the presence of cordycepin (Figure 3 A). We previously speculated that this could represent a recovery-activated mechanism involving the release of a transcriptional repressor or chromatin change (10). However, our results herein suggest that this may reflect residual transcription, based on observing steady pre-mRNA levels from late stress into recovery (Figure S1), paired with changes in transcript stability. Indeed, half-life measurements on 3,960 transcripts revealed destabilisation during light stress (median half-life decrease = 4.21 mins), which appears to facilitate RRGD. We hypothesise that 20 minutes of stress are required for this destabilisation based on expression profiling of RRGD loci: *AT3G14200*, *HSP101*, *ROF1*, and *GOLS1* (Figure S4). It remains to be established if these observations can be generalised across the transcriptome, and whether the same duration is required for different stressors.

The destabilisation of stress-induced transcripts has been observed more broadly including in yeast exposed to oxidative stress, induced by 25 minutes of hydrogen peroxide (53) and 6-30 minutes NaCl (54), and Arabidopsis cells undergoing 24 hours of cold stress (55). That stress-responsive genes are destabilised during stress appears counterintuitive. However, this likely confers faster responsivity since unstable transcripts have shorter transition times in reaching new steady state levels (12, 13). Such an inverse relationship in which unstable transcripts are more quickly induced has been observed in multiple eukaryotes, in a gene-and stress-specific manner (56, 57). While such a strategy is energetically costly, the benefits are faster responses to both the initial stress and perception of recovery (53). This response is not necessarily global, as transcript-specific responses are also observed, likely influenced by the type and duration of the stress. For example, 15 minutes of osmotic shock in yeast led to widespread transcript destabilisation, while a small subset (121 genes, 2.2%) of stress-induced transcripts were preferentially stabilised (58). In contrast, the stabilisation of stress-induced transcripts was observed in yeast experiencing a slow enduring stress (40 minutes methyl methanesulfonate treatment) (53). Similarly, in Arabidopsis, longer term salt stress (2 weeks) was also associated with the stabilisation of salt-induced transcripts encoding stress-response proteins, alongside increased N^6^-methyladenosine deposition (15, 59). This preferential stabilisation likely reflects the activity of mRNA binding proteins controlling transcript localization. For example, SPI functions to relocalize its mRNA targets towards processing bodies for stabilisation during salt stress (60).

### Rapid recovery gene down-regulation occurs as a result of stress-dependent mRNA destabilisation and, possibly, transcriptional shut-off

The rapid decline in RRGD transcript levels in recovery appears to be facilitated by their destabilisation during light stress. Indeed, previously defined RRGD loci that are highly up-regulated during light-stress were among the most destabilised transcripts (Figure 3 C). We hypothesise that transcriptional shut-off, which should occur once the stimulus for induction is removed, also contributes to this resetting although this was only verified for three RRGD loci: *HSP101*, *ROF1*, and *GOLS1*. We estimated transcriptional changes by detecting pre-mRNA through use of intronic primers amplifying intron-exon junctions of unspliced mRNA (61). Such changes are inferred under the assumption that the conversion of pre-mRNA to mRNA is constant, which could be disrupted by differential splicing or nuclear RNA decay. Unsurprisingly, changes in pre-mRNA levels tracked closely with those of mature mRNAs for the genes profiled (Figure S1). However, important differences were observed that likely reflect differential regulation of distinct light-responsive transcripts. From previous observations, *HSP101* and *ROF1* decline in abundance after 30 minutes of high light, whereas *GOLS1* declines specifically during recovery (10). For *HSP101* and *ROF1*, pre-mRNA levels (and, by proxy, transcription) declines to below pre-stress levels during recovery, effectively ‘shutting-off’ transcription, while mRNA levels remain comparatively elevated. By contrast, *GOLS1* pre-mRNA levels more closely resembled those in mRNA, and did not decline to or below pre-stress levels during recovery. This likely reflects differing transcriptional repression of light-responsive transcripts during recovery, which could be further unpacked with more extensive profiling of transcriptional changes. This will be most practical using genome-wide measurements of pre-mRNA levels that may be possible from conventional mRNA sequencing at greater sequence depth (62), since the short timescales and stress treatments used here make protocols such as base analog labelling, RNA polymerase II ChIP sequencing, and global run-on sequencing impractical. Notably, however, pre-mRNA levels began decreasing during the stress period for all three genes assayed, regardless of changes in mRNA. This suggests an initial transcriptional burst, mediated by light-activated transcription factors, which lasts for approximately 30 minutes as observed in mammals and fungi in response to a stimulus (63–65).

Transcriptome resetting during recovery appears to be determined in advance, during light stress, by changes in mRNA stability and transcription. As these are influenced by the duration of stress, it stands to reason that the recovery response may be altered in the same way. Among studies investigating recovery, it is rare that differing lengths of stress are compared. As demonstrated here, the recovery response is influenced by the duration of stress. Indeed, plants entering recovery after only ten minutes of light stress displayed steady transcript abundance (Figure S4). Plants required at least 30 minutes of light stress before observing the expected rapid decline, which coincides with the mRNA stability data showing transcripts were destabilised by this point. Given that transcription appears to be ongoing at 30 minutes, this suggests that transcript destabilisation is the primary response for regulation in recovery, with a secondary transcriptional shut-off occurring during recovery itself. This coordination of mRNA stability and transcription aligns with observations made in yeast responding to oxidative stress (53, 54), whereby early induced transcripts are destabilised while those repressed are stabilised. This appears to be a mechanism by which the organism ‘prepares’ for recovery; upon which the destabilised stress-induced transcripts are degraded while the stabilised stress-repressed transcripts accumulate, ultimately resetting the transcriptome to a pre-stress state.

Impairments in RNA decay enzymes, especially 5′ - 3′ RNA degradation pathways, are well documented in influencing plant stress responses (17–23, 25). However, the mechanism of transcript destabilisation, occurring during light stress, which facilitates RRGD remains unclear. Indeed, this has been challenging to identify since RNA decay pathways are complex and redundant, with mutant analyses complicated by both feedback, compensation, and lethality of combinatorial mutations. Previous studies have profiled a series of combinatorial mutants during recovery, however, down-regulation of stress-induced genes appeared unperturbed (10, 66). mRNA turnover is also regulated by numerous mRNA binding proteins, interacting with translation and decay machinery, which require further exploration (40). Post-translational regulation of the activity of RNA helicases, which degrade stress-responsive transcripts, presents one possible mechanism (43). Another is the involvement of stress-induced variation in 5′-end NAD^+^ capping for preferential transcript-specific changes in stability (67).

### Regulation of translation during late light stress and at onset of recovery

There is an increasing number of reports highlighting the regulation of translation in plants responding to stress, at both global and transcript-specific levels in conjunction with changes in mRNA turnover and stability (40). Extending to this literature, we find that regulation of translation is dynamic during light stress and recovery. This was evident as transcript-specific changes in polysome-association (Figure S5 A-B), as opposed to wholesale reorganisation elicited by pattern-triggered immunity (bacterial translation elongation factor EF-Tu, elf18) (68), darkness (69), heat stress (32, 70), and hypoxia (71). The modesty in translational response observed here may reflect a stress of lesser intensity, in this case a 10-fold increase in light irradiance for 60 minutes. This posits greater complexity to the regulation of translation, at the global and transcript-specific levels, to be tuned based on environmental stimuli and requirements. This is likely determined through the action of intra-cellular signalling pathways and their interplay with RNA-binding proteins and translational machinery, which represents new avenues of investigation. Retrograde signalling has been linked to nuclear RNA processing (72). However, the speed with which translational regulation occurs (e.g. within 7.5 minutes recovery, Figure S5 C) posits direct interaction between chloroplast-derived signals and cytosolic translation factors or interacting RNA-binding proteins, many of which are redox-sensitive (73, 74).

Temporal differences in transcript-specific translation were clearly observable based on changes in RPL (Figure 4 B). During early stress, there was a slight bias toward decreased RPL, indicating translation repression of many genes. This aligns with observations that stress typically initially represses translation by perturbing translational machinery involved in initiation, scanning, and elongation (30, 39, 75). On the other hand, we observed a striking increase in RPL during late stress that occurred independently of changes in total mRNA levels. This likely reflects the combination of two phenomena. In the first instance, this may reflect a transient pausing of elongation, without eliciting mRNA degradation, which resumes as cells begin to acclimate to the new conditions, as was observed in yeast (75). Simultaneously, preferential translation may be occurring for light stress-associated genes to facilitate an acclimatory response (34). The concordance between total and polysome-bound mRNA levels was also uncoupled rapidly upon transition to recovery, but is eventually re-established after 30 minutes (Figure S5 C). This reinforces the notion that this is an active period of regulation to establish post-stress homeostasis to favour growth, as opposed to a passive release of stress defence to return to a pre-stress condition. Indeed, although physiological and anatomical parameters return to pre-stressed levels after recovery from non-lethal stress, differential redox (76), proteomic (77–79), and metabolomic (80, 81) signatures represent a unique cellular environment especially during early recovery.

Translation and mRNA stability are coupled by a complex variety of interactions (26). For instance, perturbations to translation, such as ribosome initiation, elongation, or unloading, can elicit changes to transcript stability in association with mRNA translation and degradation machinery. For instance, glucose withdrawal rapidly abolished 5’ RNA binding of translation initiation factors within 2 minutes, leading to translational shutdown without mRNA degradation (39). Conversely, heat shock induced progressive loss of 5’ RNA binding of translation initiation factors paired with XRN1-mediated mRNA degradation within 12 minutes (39). Given the complexity of these interactions, it is unsurprising that we observed a negligible relationship between changes in RPL and stability (Figure 4 C). However, subsets of transcripts showed changes in polysome-association that were consistent with protection from degradation and co-translation decay. For example, many transcripts had altered RPL without significant changes in total mRNA abundance (e.g. 13.22% after 30 mins HL, Figure 4 B). At the same time, a smaller fraction (6.64% after 30 mins HL) of transcripts showed increased or unchanged RPL alongside decreased total mRNA abundance, possibly representing transcripts undergoing co-translational decay. Contrary to expectations, we did not see evidence of elevated co-translation decay during recovery, with a similar proportion of transcripts having reduced mRNA abundance alongside unchanged, or increased, RPL (5.43%). Further, it was hypothesised that RRGD transcripts underwent co-translational decay during recovery, which we expected to observe as reduced RPL either during late stress or upon recovery. While declines in total mRNA of RRGD loci were generally concordant with declines in polysome-bound mRNA (Figure S6), RPL tended to increase during late stress in high light induced transcripts (Figure 5). Upon recovery, there was a notable decrease in RPL, especially for category 3 RRGD transcripts, suggesting that polysome-bound mRNAs were declining faster than free mRNAs, and therefore are likely undergoing co-translational decay. The incorporation of translational inhibitors with this experimental strategy, targeting initiation (82) or elongation (83), may help unpack the fate of light-induced transcripts when translation is perturbed.

Lastly, it was previously unclear the extent to which rapid changes in mRNA abundance would contribute to protein-level changes (10). In general, we found that changes in polysome-bound mRNA levels correlated with those in the total mRNA pool. This was highlighted by temporal changes measured in previously defined RRGD loci (Figure 5, Figure S6). Therefore, transcript-level changes in total mRNA should translate to increased ribosome association, resulting in greater translation. Whether or not this would lead to gross changes in protein abundance is unclear, depending on the stability of the protein product (51). Nonetheless, our observations suggest that the transcriptional compensation observed at genes encoding proteins degraded under high light, should result in elevated translation that contributes to maintaining proteostasis during stress (84).

## Materials and Methods

### Plant growth and treatments

Arabidopsis seeds were sown onto moist soil (Martins Seed Raising and Cutting Mix) supplemented with Osmocote Exact Mini slow release fertiliser (Scotts) at 1 g/L dry volume of soil and 1 L of 0.3 % (v/v) AzaMax (Organic Crop Protectants). Seeds were covered with plastic wrap and stratified at 4 °C in the dark for at least 72 hours to break dormancy and synchronise germination. Stratified seeds were transferred to a temperature controlled Conviron S10H growth chamber (Conviron, Canada) fitted with a mixture of 250 W metal halide (Venture Lighting, MH 250W/U) and high pressure sodium lamps (Phillips, SON□T 250W E E40 SL/12). Plants were cultivated under a 12-hour photoperiod of 100 μmol photons m^-2^ s ^-1^, 21 °C, and 55 % relative humidity. Plants were watered to saturation the day preceding light stress application. Light stress was induced by increasing the light intensity 10-fold (i.e. 1000 μmol photons m^-2^ s ^-1^), resulting in a “warm” high light treatment (simulating sunlight, ΔT = 7 °C) that effectively induces oxidative stress (10, 85). For recovery, plants were returned to pre-stress light conditions. Plants were moved between pre-programmed growth chambers to impose light stress and recovery. All experiments were performed at the midday point of the light period to exclude diurnal effects.

For transcriptional inhibition, cordycepin (3’-deoxyadenosine, Sigma-Aldrich) was syringe infiltrated on the abaxial side of fully expanded leaves (true leaves 4-6) of 28-day old plants. Individual leaves were infiltrated with approximately 100 µL of incubation buffer [1 mM PIPES (pH 6.25), 1 mM sodium citrate, 1 mM KCl, 15 mM sucrose] with 0.6 mM cordycepin (86, 87), or without (mock), using a 1 mL needleless syringe (Terumo). Once the buffer had permeated through the entire leaf, the syringe was removed and plants were pre-incubated for 10 minutes (unless otherwise stated) before further treatment and harvesting. Each biological replicate represents leaves from independently grown plants infiltrated with either mock or cordycepin-containing buffer. Paired leaves were sampled from the same plant at each time point per condition: unstressed (US), high light (HL), or recovery (REC). Infiltrated leaves were excised, at the appropriate time point under each condition, from the base of the petiole and immediately flash-frozen, in a 2 mL safe-lock microcentrifuge tube (Eppendorf), using liquid nitrogen. Frozen tissue was stored at -80 °C until ready for processing.

### RNA isolation

Frozen tissue was ground into a fine powder using a ⅛” steel ball bearing with 1 min shaking at 25 Hz in a TissueLyser II (Qiagen). Total RNA was extracted from finely ground tissue using TRI reagent (Sigma-Aldrich, #T9424-200ML) at a ratio of 1 mL solution per 100 mg ground tissue. Residual phenol was removed from the crude extract through two chloroform extractions at a ratio of 1:5 (v/v), followed by precipitation using isopropanol at 1:1 (v/v). Precipitated RNA was washed twice with 70% ethanol and resuspended in 1 mM sodium citrate buffer (pH 5.4). RNA quantification was performed through spectrophotometric analysis at 260 nm using the Nanodrop ND-1000 Spectrophotometer and RNA quality was assessed using 1% agarose gel electrophoresis or the LabChip GX Touch (PerkinElmer). 5 µg of purified total RNA was combined with 5 µL of TURBO DNase buffer and 1 µL TURBO DNase (Thermo Fisher Scientific) in a 50 µL reaction, and incubated at 37°C for 30 minutes. DNA nuclease-treated RNA was then purified using 1.8□ Sera-Mag paramagnetic particles.

### qRT-PCR analysis

DNA nuclease-treated RNA was reverse transcribed into cDNA using either Invitrogen Superscript III (Thermo Fisher Scientific) or Maxima H Minus (Thermo Fisher Scientific) reverse transcriptase according to the manufacturer’s instructions. For detection of mRNAs, 1 µg of DNA nuclease-treated RNA was combined with dNTPs and an Oligo(dT)_18_ primer in a 4.5 µL reaction volume to final concentrations of 2.2 mM and 11.1 µM, respectively, and incubated for 5 mins at 65°C. For Superscript III based reverse transcription, 1□ first strand reaction buffer, 10 mM DTT and 100 units of Superscript III Reverse Transcriptase was added to a final volume of 10 µL before incubation at 50°C for 60 minutes and 70°C for 15 minutes. For Maxima H Minus based reverse transcription, 1□ reverse transcription buffer and 100 units Maxima H Minus Reverse Transcriptase was added to the reaction mix before incubation at 50°C for 30 minutes and 80°C for 5 minutes. For the detection of pre-mRNA, 500 ng of DNA nuclease-treated RNA was combined with 1.05 mM dNTPs and 10.5 µM random hexamers (Qiagen) in a reaction volume of 14.25 µL before incubation at 65°C for 5 minutes. 1 first strand reaction buffer, 5 mM DTT and 150 U Superscript III (Thermo Fisher Scientific) were added to a final volume of 14.25 µL, before incubation at 25°C for 10 minutes, 50°C for one hour, and 70°C for 15 minutes. Expression of mRNA and pre-mRNA was then assayed by semi-quantitative reverse-transcription PCR (qRT-PCR) on a Roche LightCycler480 using SYBR Green I (Roche Diagnostics). Raw fluorescence data were analysed using LinRegPCR to perform background subtraction, determine PCR efficiency, and calculate starting concentration [N_0_; arbitrary fluorescence units (88, 89)]. N_0_ values were used to calculate fold-changes for target genes, which were normalised to changes in the housekeeper *PROTEIN PHOSPHATASE 2A SUBUNIT A3 (PP2AA3, AT1G13320)*. At least three biological replicates per genotype per time point were sampled, and each qPCR reaction was run in technical duplicate or triplicate. Primer sequences are provided in supplementary table 1.

### mRNA sequencing

The Illumina TruSeq Stranded mRNA kit was used to prepare strand-specific (reverse-stranded: first read maps to the reverse strand) polyA-enriched sequencing libraries according to the manufacturer’s instructions, except that all reactions were scaled-down by one-third and SuperScript III (Invitrogen) was used for first strand synthesis at 50 °C. Libraries were constructed using Illumina unique dual index adapters in a 15 cycle indexing PCR. All clean-up steps were performed using nuclease-free AMPure XP beads (Beckman Coulter) or Sera-mag SpeedBeads (GE Healthcare). Library concentration and fragment distribution were determined on the Qubit (double-stranded DNA high sensitivity kit, Invitrogen) and LabChip GX-Touch (DNA High Sensitivity kit, PerkinElmer), respectively. The concentration and peak fragment size were used to determine individual library molarity (assuming the average molar mass of one DNA bp = 650 g/mol). Individual libraries were pooled in equal molar ratios and sequenced on a NextSeq500 (75 bp single-end, supplementary table 2) at the ACRF Biomolecular Research Facility (Australian National University).

**Table 2.**
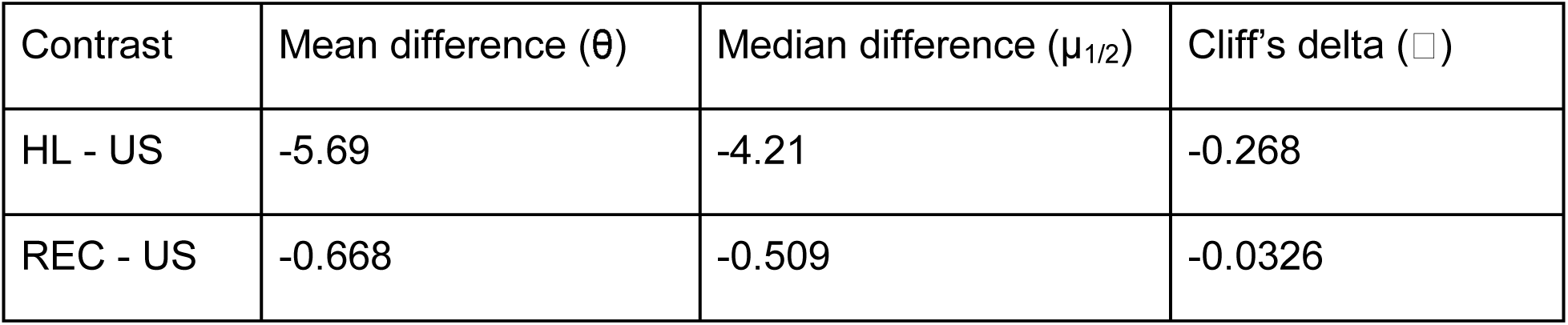
Effect sizes parameters for differences in transcript half-life between conditions

| Contrast | Mean difference ($\theta$) | Median difference ($\mu_{1/2}$) | Cliff's delta (□) |
| --- | --- | --- | --- |
| HL - US | -5.69 | -4.21 | -0.268 |
| REC - US | -0.668 | -0.509 | -0.0326 |

Raw sequencing reads were trimmed to remove adapter sequences and low-quality base calls (PHRED < 20, -q 20) with *Trim Galore!* (v0.6.4), a wrapper for *Cutadapt* (v1.18), followed by inspection with *FastQC* (v0.11.8). Trimmed reads were used for transcript quantification using *Kallisto* (90). *Kallisto index* (-k 21) was used to build a transcript index based on TAIR10 coding sequences [Ensembl release 51: Arabidopsis_thaliana.TAIR10.cdna.all.fa.gz (91)]. Bootstrapped transcript-level abundances were computed using *Kallisto quant* (--rf-stranded --bias --single -b 10 -l 300 -s 100).

Transcript-level abundance estimates were summarised to gene-level counts, using *Tximport* [‘lengthScaledTPM’ (92)], for genes detected at an abundance of at least 0.5 TPM in more than six *t_10_* samples (17,256 loci, supplementary table 3). The *RUVr* procedure of the *RUVSeq* package was applied to account for variation between experimental batches (93). Briefly, deviance residuals were quantified using a first-pass generalised linear model of rounded counts, without global scaling, on the covariates of interest. Then, a factor analysis is performed using the deviance residuals to compute batch-corrected counts using the *RUVr* function. Multidimensional scaling was performed on the batch-corrected counts using *plotMDS* (gene.selection = “pairwise”). Differential gene expression analysis was performed on batch-corrected counts using the *edgeR* quasi-likelihood pipeline without TMM normalisation (94). A quasi-likelihood negative binomial generalised log-linear model was fit to the adjusted counts (glmQLFit, robust=T) followed by application of quasi-likelihood F-tests (glmQLFTest), with FDR correction for multiple hypothesis testing, to detect significant differentially expressed genes (FDR adjusted p-value < 0.05).

Adjusted counts were converted to abundance (counts per million, CPM) and used to compute fractional decreases in mRNA abundance (relative to the mean at *t*_10_ per condition). The *t*_10_ time point was shared between HL and REC samples as the same batch of infiltrated plants were either retained under HL or transferred to standard growth conditions for REC. Decay factor normalisation was performed to account for the apparent increase in abundance of relatively stable genes, which constitute a greater proportion of the total RNA pool while those less stable degrade without replacement during transcriptional inhibition (44). To do this, a decay factor was computed for each time point per condition, across the cordycepin treated samples, using a set of highly abundant and stable reference genes. Reference genes were selected based on their abundance [>95th percentile (log_2_ CPM+0.01)] and variance [<25th percentile (coefficient of variation CPM)] across mock-infiltrated samples. This produced the following 52 reference genes: *AT4G01050, AT4G24280, AT5G36790, AT1G07920, AT5G13650, AT1G05850, AT1G62750, AT4G22890, AT3G62030, AT3G19170, AT2G18960, AT3G23400, AT5G42270, AT3G63140, AT3G11630, AT4G24770, AT1G65960, AT3G54050, AT4G01150, AT5G61410, AT1G11860, AT4G32260, AT5G60600, AT5G42980, AT3G55800, AT3G46780, AT3G16140, AT4G20360, AT3G14420, AT1G56070, AT5G66190, AT3G02470, AT1G30380, AT4G03280, AT2G30950, AT5G60390, AT4G04640, AT1G55670, AT3G60750, AT1G44575, AT5G35630, AT4G28750, AT1G20020, AT3G12780, AT5G50920, AT1G42970, AT1G20340, AT4G38970, AT4G37930, AT4G12800, AT5G01530, AT3G61470*. The mean fold-increase computed across reference genes in cordycepin infiltrated samples should reflect the fold-change in the total RNA pool, thus providing a scaling factor that was applied on fractional decreases at each time point per condition. Lastly, any genes still displaying a fold-increase ≥1.5 after *t*_10_ were excluded from half-life analysis leaving 14,386 and 9,339 loci for unstressed and all conditions, respectively.

### mRNA half-life analysis

Decay factor normalised fractional decreases in mRNA abundance were used to compute half-life (*t_1/2_*) per condition (assuming exponential decay and first order kinetics) (13). The decay constant (*k_d_*) was first computed using log-linear modelling of the change in mRNA abundance as a function of time using individual data points from cordycepin infiltrated samples in each condition: *k_d_ = - log_e_[C/C_10_]/dT*. From this, the condition-specific half-life of each gene could be solved using: *t_1/2_ = log_e_(2)/k_d_*. The *R* package *lme4* was used for building linear mixed-effects models with fixed (Time) and random variables (experimental batch) (95). The conditional *R^2^* and accompanying F-statistic and p-value were used to evaluate whether the data fit an exponential decay model (p-value < 0.05) on a gene-by-gene basis, with the *R* packages *piecewiseSEM* and *lmerTest* (96, 97). Finally, any genes calculated with *k_d_* < 0 or half-life > 1,440 min (1 day) were removed from the analysis. In total, 6,711 and 3,960 genes could be statistically modelled under unstressed and all conditions, respectively. Half-lives calculated under unstressed conditions were directly compared to those reported for a common set of 4,497 genes across multiple genome-wide studies in Arabidopsis (11, 41, 43, 44). The R package *dabestr* was used to compute paired effect size parameters: paired mean difference (θ), paired median difference (µ_1/2_), and paired Cliff’s delta (□) using the functions *mean_diff*, *median_diff*, and *cliffs_delta*, respectively, with 10,000 bootstrap resamples. Significant functional enrichments (FDR < 0.05) were assessed using *ShinyGO v0.76.1* (98).

### Polysome-associated mRNA sequencing

Polysome-associated mRNA sequencing was performed using an adapted protocol (99) with an isolation buffer of lower ionic strength, based on buffer D from (46), with greater yields of intact ribosome from plant tissue. Briefly, 250 mg of ground plant tissue was dissolved in 1 mL of polysome extraction buffer [160 mM Tris-Cl (pH 7.6), 80 mM KCl, 5 mM MgCl_2_, 5.36 mM EGTA (pH 8), 0.5% IGEPAL CA-630, 40 U/mL RNasin Plus RNase inhibitor (Promega), 150 µg/mL cycloheximide, and 150 µg/mL chloramphenicol] and incubated on ice for 10 minutes. Samples were repeatedly centrifuged at 16,000 rcf for 5 minutes at 4°C until the supernatant was clear. Samples were loaded onto sucrose gradients, consisting of layers of 50% (1.68 mL), 35% (3.32 mL), 20% (3.32 mL) and 20% (1.68 mL), with the two 20% layers added separately (99). The buffer used in the gradients consisted of 400 mM Tris-Cl (pH 8.4), 200 mM KCl, 100 mM MgCl_2_, 10.12 µg/mL cycloheximide and 10.12 µg/mL chloramphenicol. Gradients were centrifuged at 41,000 rpm with an SW41Ti rotor for 2 hours at 4°C. Each gradient was passed through a spectrophotometer from highest to lowest density and absorbance at 260 nm recorded, with 1 mL fractions collected. Fractions were pooled into monosomal and polysome sets and extracted using 5:1 acid phenol:chloroform. Briefy, an equal volume of the phenol:chloroform mix was added to each fraction, before centrifugation at 16,000 rcf for 10 minutes. The aqueous layer was extracted a second time using a one-fifth volume of chloroform, before the RNA was precipitated using an equal volume of isopropanol in addition to sodium acetate (pH 5.2) to a final concentration of 300 mM. The precipitated RNA was washed twice with 70% ethanol and resuspended in a 1 mM sodium citrate (pH 5.4) solution. Purified polysome RNA, and total mRNA purified from paired ground tissue (purified as described above), were used to generate sequencing libraries using the Illumina TruSeq Stranded mRNA kit as described above.

Quality control of raw reads was carried out using *FastQC*, with adapter trimming performed using *scythe* (-p 0.1) and quality trimming performed using *sickle* (-q 20 -l 20). Reads were aligned to the TAIR10 Arabidopsis genome using *subjunc* to report a single unambiguous mapping location per read (100). Sorting, indexing, and compression was carried out with samtools (101) and read counts per loci were calculated using *featureCounts* [-s 2 for reverse stranded libraries, (102)]. Polysome-bound mRNA samples in replicate one were omitted from the following analysis due to limited read depth. Reads mapping to rRNA were removed before performing TMM normalisation and differential gene expression analysis, and calculating reads per kilobase of transcript per million mapped reads (RPKM) at the gene-level in *edgeR* (103). Quasi-likelihood F-tests were performed, as described above, to detect significantly differentially ribosome-associated transcripts. Relative polysome loading (RPL) was calculated by dividing the RPKM of polysome-bound RNA by that of total RNA on a per replicate basis.

### Data availability

All code used for analyses are available on GitHub (https://github.com/dtrain16/NGS-scripts). All sequencing data are accessible at the NCBI’s Gene Expression Omnibus (104) at accession number GSE201015 (https://www.ncbi.nlm.nih.gov/geo/query/acc.cgi?acc=GSE201015).

## Author contributions

PAC and BJP conceived the project. ABS, DRG, MM, BJP, and PAC designed experiments. ABS and DRG performed *in planta* cordycepin experiments and mRNA sequencing. ABS, MM, YJ, and NS performed polysome profiling. ABS, DRG, and AFB performed data analysis. ABS, DRG, and PAC led manuscript preparation. All authors read and commented on the manuscript.

## Acknowledgements

This work was funded by the Australian Research Council (DE200101748 to PAC, FL190100056 to BJP, DP220103640 to MM, DRG, and BJP), the CSIRO Synthetic Biology Future Science Platform (to DRG), the Grains Research and Development Council (GRS11010 to ABS), and the Australian Government Research Training Program (to ABS). We acknowledge the ACRF Biomolecular Resource Facility, a service node of Bioplatforms Australia, and the Centre for Biodiversity Analysis Ecogenomics and Bioinformatics Lab, both hosted at The Australian National University, for provision of resources and expertise to perform Illumina sequencing. This project was supported with the provision of plant growth facilities by the Australian Plant Phenomics Facility and computational infrastructure by the National Computational Infrastructure, both supported under the National Collaborative Research Infrastructure Strategy of the Australian Government.

---

**Figure S1.**
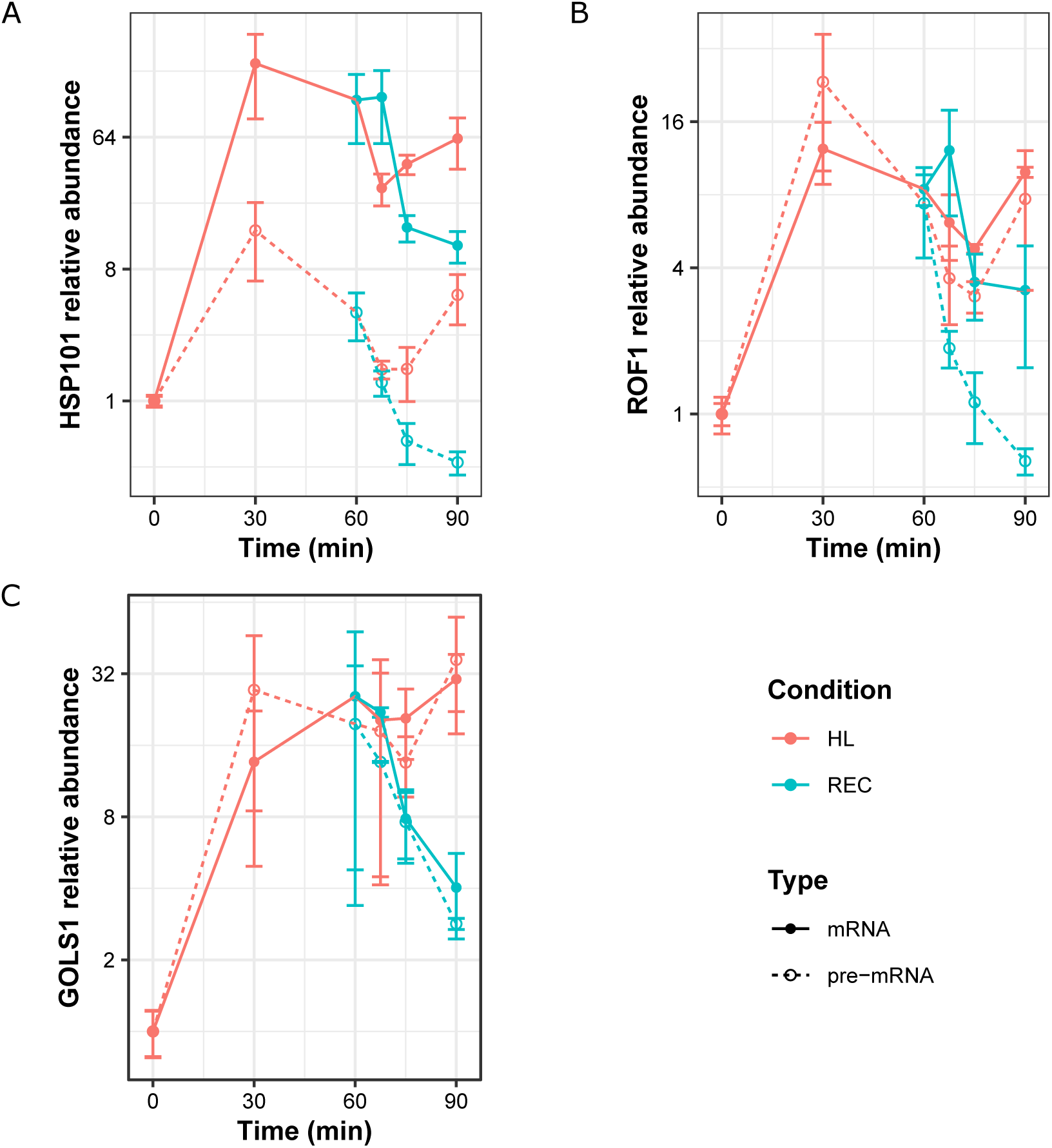
Pre-mRNA levels decrease more rapidly during recovery than mRNA. (A-C) Profiles of pre-mRNA and mRNA for *HSP101*, *ROF1*, and *GOLS1* during HL and REC. Data is presented as relative abundance compared to time 0. Points denote means, error bars denote standard error of the mean (n=3).

**Figure S2.**
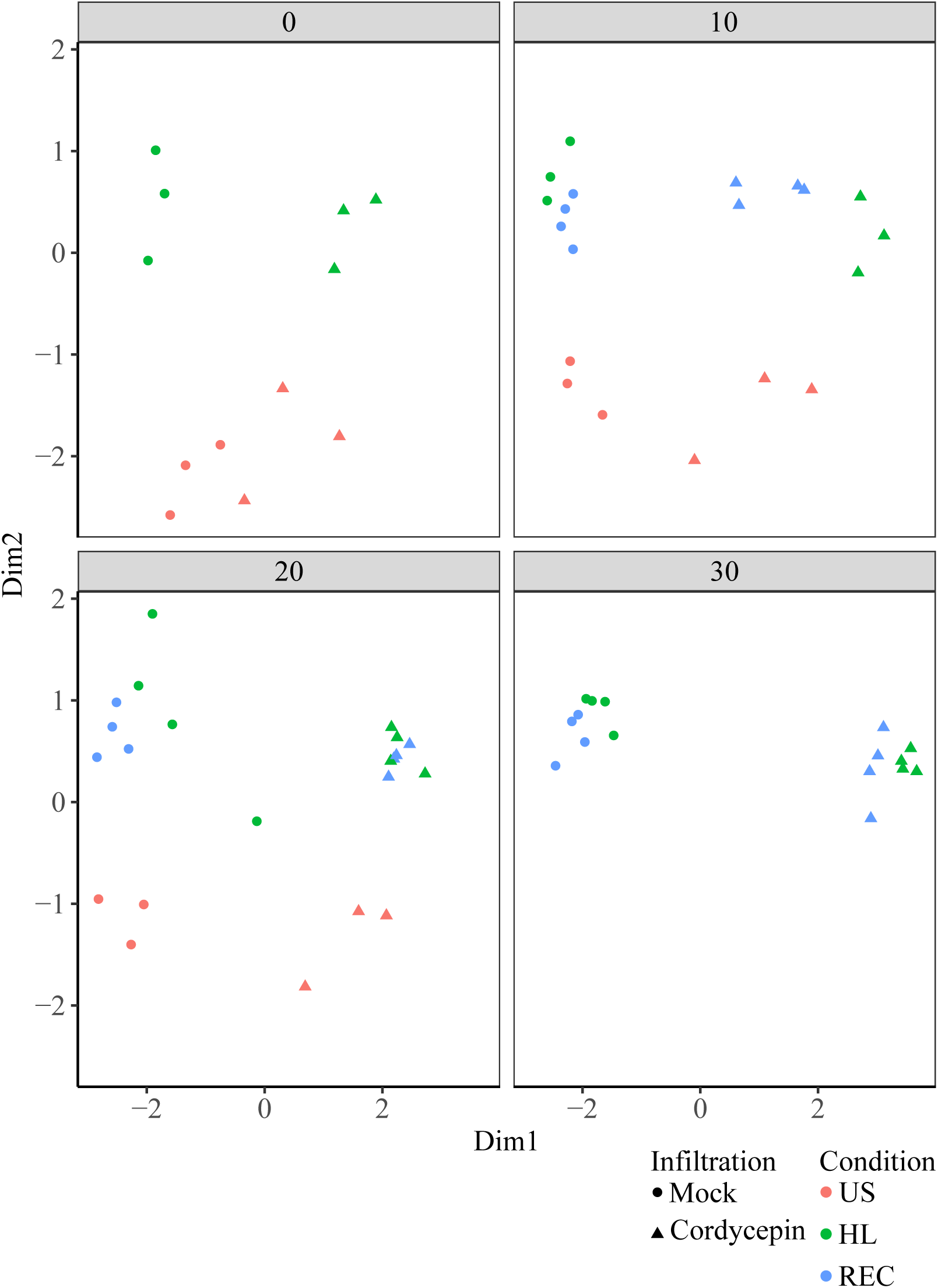
Cordycepin timecourse sample clustering. Multi-dimensional scaling plot of all samples where distance reflects the typical log_2_ fold-change between samples for the genes that distinguish those samples. Point shapes denote infiltration (mock or cordycepin) and colour denotes condition (US, HL, or REC).

**Figure S3.**
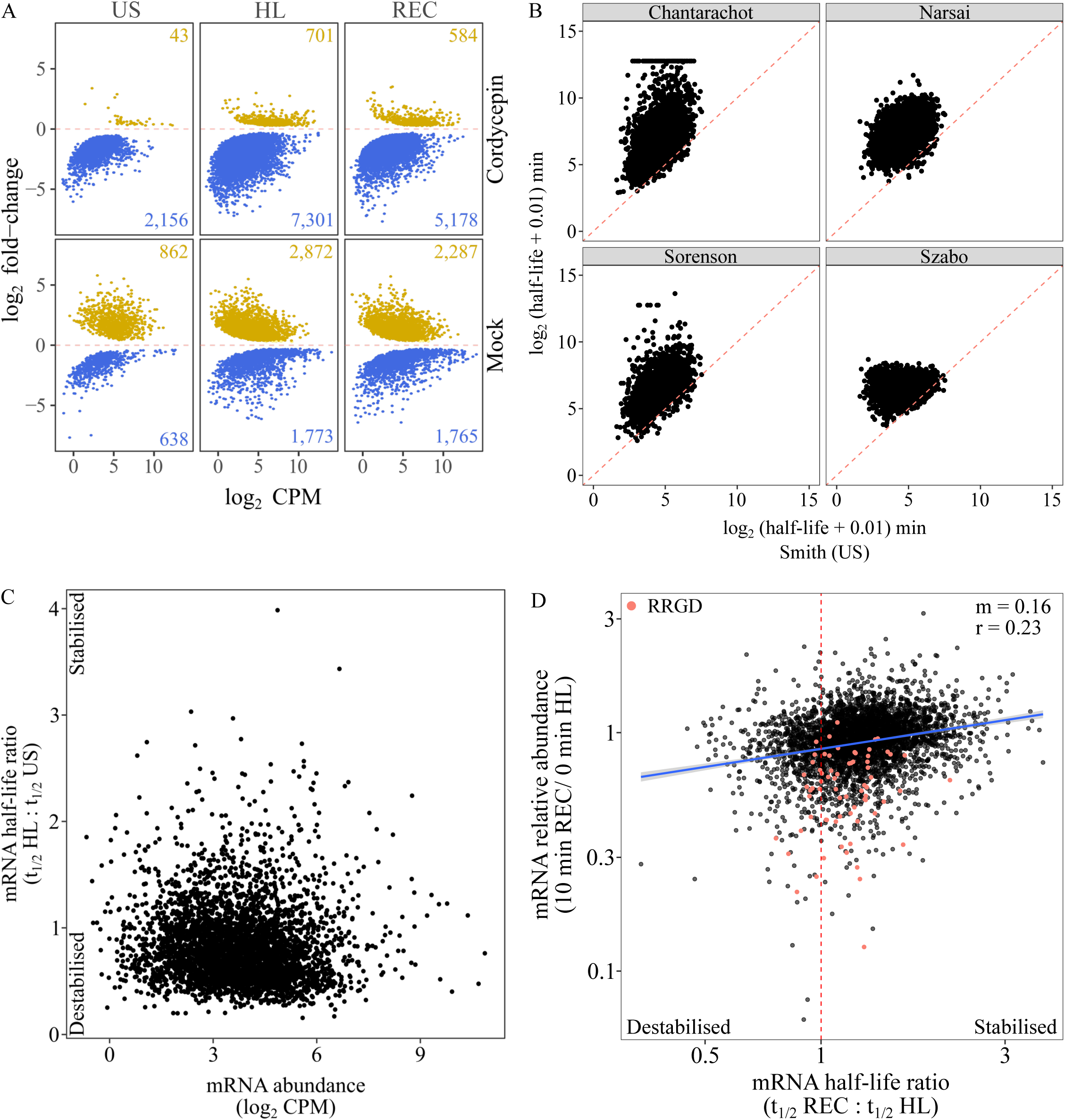
Syringe-infiltration of cordycepin facilitates modelling half-lives for transcripts of varying abundance. (A) Differential gene expression between the first and last time points under cordycepin or mock treatment in US, HL, or REC. Up-and down-regulated genes (p < 0.05) are denoted by yellow and blue points, respectively. (B) Scatter plots comparing transcript-specific half-lives (log_2_ halflife + 0.01) for the 3,960 modelled in this study (x-axis) compared to prior work (y-axis). (C) Scatter plot relating the half-life ratio (HL/US) to mean mRNA abundance, calculated across mock treated US and HL samples at 10 minutes, for the 3,960 modelled transcripts. Points represent individual genes. (D) Scatter plot presenting the fold-change in mRNA abundance (10 min REC/ 0 min HL), in mock-treated samples, versus the half-life ratio (REC/HL) determined for 3,960 modelled transcripts. Points denote individual genes, line denotes fitted linear model, ‘m’ denotes regression coefficient, and ‘r’ denotes Pearson’s correlation coefficient.

**Figure S4.**
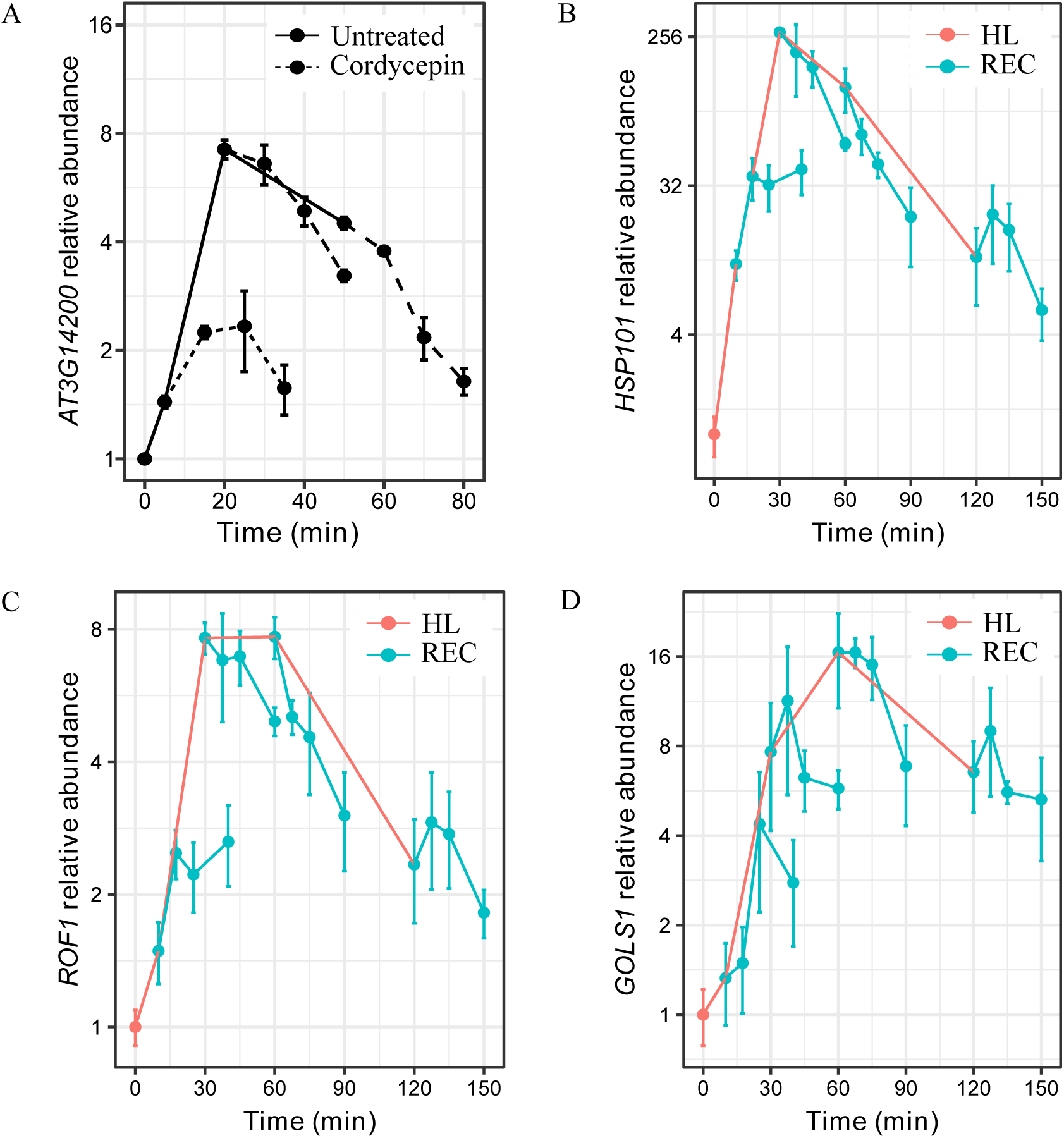
Rapid recovery gene down-regulation is dependent on length of HL. (A) Gene expression changes in AT3G14200 following cordycepin infiltration carried out at 5, 20, and 50 minutes HL. Solid line denotes untreated leaves, dashed line denotes cordycepin-infiltrated samples. (B-D) Gene expression profiles of *HSP101*, *ROF1*, and *GOLS1* during REC after various lengths of HL. Red lines denote HL, blue lines denote REC (initiated at 10, 30, 60, and 120 minutes). Points denote means, error bars denote standard error of the mean (n = 3).

**Figure S5.**
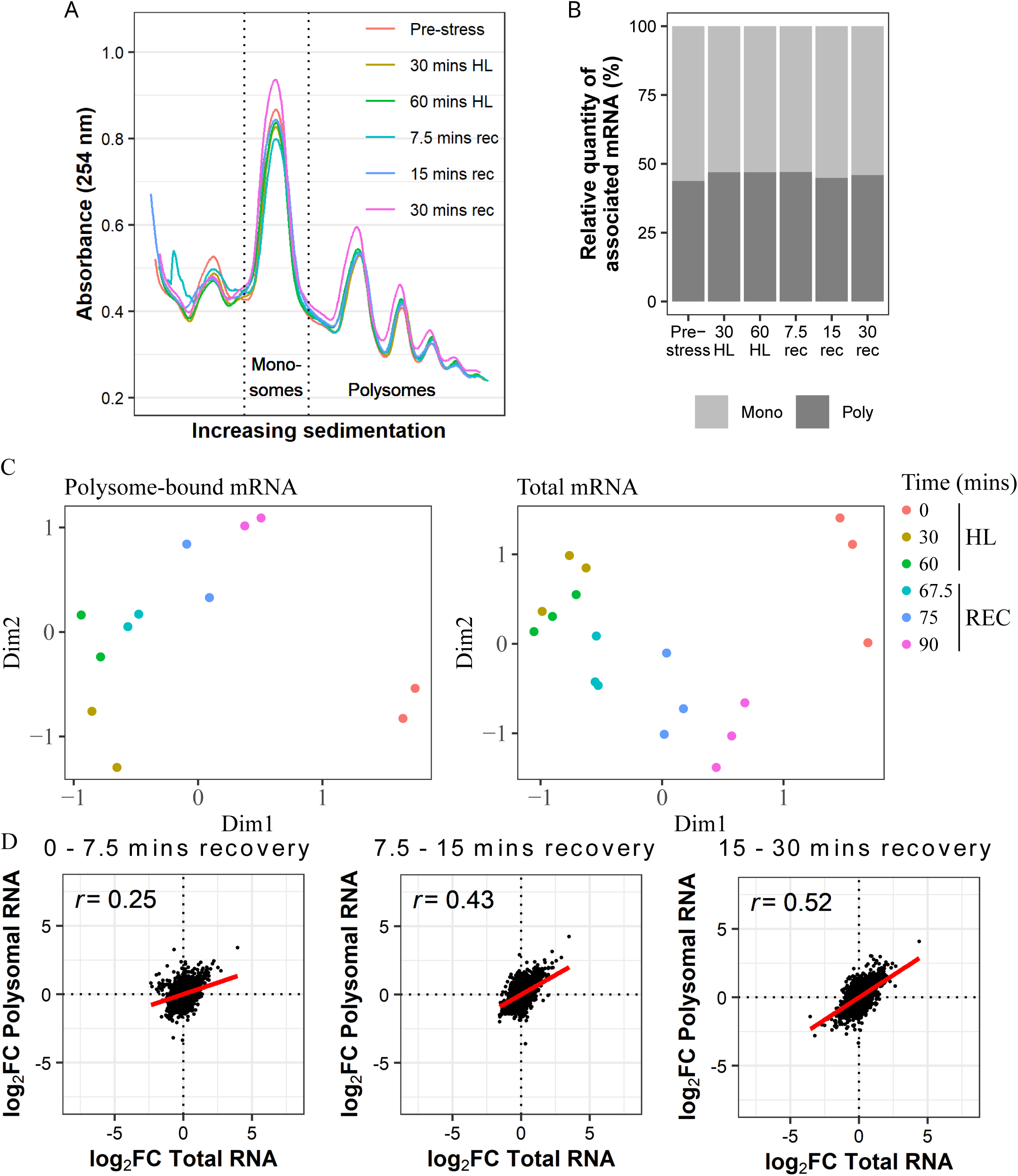
Uncoupling of total and polysome-bound RNA abundance during early recovery. (A) Monosomes and polysomes extracted from Arabidopsis seedlings were separated using gradient centrifugation and quantified by absorbance at 254 nm. (B) Relative abundance of monosomes and polysomes was calculated by integration of the area under the absorbance profile. (C) Multi-dimensional scaling plots of polysome-bound and total mRNA sequencing samples, during a light stress and recovery timcourse. Distance reflects the typical log2 fold-change between samples for the genes that distinguish those samples. (D) Correlation between changes in total and polysome-bound mRNA abundance between recovery time points. r denotes Pearson’s correlation coefficient.

**Figure S6.**
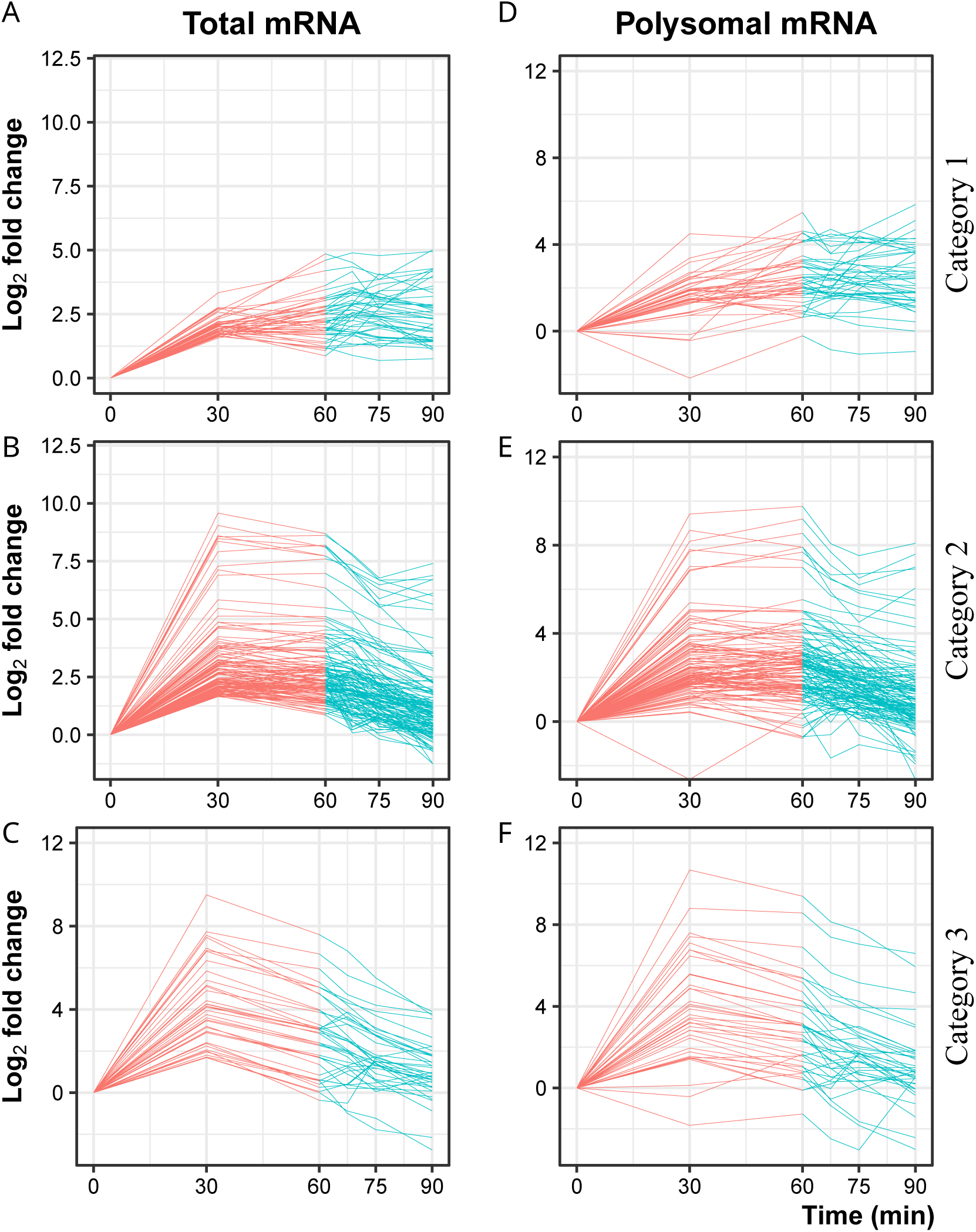
Expression profiles of light-induced total and polysome-associated mRNAs during light stress and recovery. (A-C) Genes induced by HL (FC > 3 after 30 mins) were categorised by their expression profiles in the total mRNA fraction. Category 1 genes remained elevated during REC. Category 2 genes were significantly down-regulated during REC. Category 3 genes were significantly down-regulated between 30 and 60 minutes HL. (D-F) The corresponding changes in polysome-bound mRNA are displayed. Lines represent mean relative abundance over time during HL (red) and REC (blue).

